# Computational design of BclxL inhibitors that target transmembrane domain interactions

**DOI:** 10.1101/2022.11.09.515782

**Authors:** Gerard Duart, Assaf Elazar, Jonathan J. Weinstein, Laura Gadea-Salom, Juan Ortiz-Mateu, Sarel J. Fleishman, Ismael Mingarro, Luis Martinez-Gil

**Affiliations:** Department of Biochemistry and Molecular Biology, Institut de Biotecnologia i Biomedicina BIOTECMED, Universitat de València, 46100, Burjassot, Spain; Department of Biomolecular Sciences, Weizmann Institute of Science, Rehovot, Israel

## Abstract

Several methods have been developed to explore interactions among water-soluble proteins or regions of proteins. However, techniques to target transmembrane domains have not been examined thoroughly. Here we developed a novel computational approach to design transmembrane sequences that specifically modulate protein-protein interactions in the membrane. To illustrate this method we demonstrated that BclxL can interact with other members of the Bcl2 family through the transmembrane domain and that these interactions are necessary for BclxL control of cell death. Next, we designed sequences that specifically recognize and sequester the transmembrane domain of BclxL. Hence, we were able to prevent BclxL intra-membrane interactions and cancel its anti-apoptotic effect. These results advance our understanding of protein-protein interactions in membranes and provide new means to modulate them. Moreover, the success of our approach may trigger the development of a new generation of inhibitors targeting interactions between transmembrane domains.

## Introduction

Many methods, including computational approaches(1–3), have been developed to explore interactions among water-soluble proteins or regions of proteins. However, techniques to target transmembrane domains (TMDs, TMD singular), despite their importance in a wide variety of cellular processes, have not been examined thoroughly because of the intrinsic difficulty of working with membrane proteins and lack of understanding of TMD–TMD interactions. In a seminal work, Yin et al. described a method for the computational design of peptides that target TMDs in a sequence-specific manner(4). To illustrate their method, the authors designed and tested peptides that recognized the TMD of two integrins *in vitro* and *in vivo*. In more recent work, Elazar et al. designed single-pass α-helical TMDs that self-assembled, basing their approach on an improved *ab initio* Rosetta atomistic modeling strategy(5–7). These TMDs were incorporated into a chimeric antigen receptor (CAR), and results indicated that *in vitro* CAR T-cell cytokine release and *in vivo* anti-tumor activity scaled linearly with the oligomeric state encoded by the receptor TMD(8).

Apoptosis, the main mechanism of controlled cell death, is a conserved cellular process that occurs in response to various physiological and pathological situations(9). There are two signaling routes, the so-called extrinsic and intrinsic pathways, that lead to apoptosis. The intrinsic pathway begins with the release of apoptogenic factors from the mitochondria which in turn prompts caspase 3 activation and ultimately apoptosis. Permeabilization of the mitochondrial outer membrane (MOM) is primarily regulated by the B-cell lymphoma 2 (Bcl2) protein family(10). In healthy cells, anti-apoptotic Bcl2 members inhibit the activation of pro-apoptotic proteins through direct interaction or by sequestering BH3-only proteins(11–14). Upon an apoptotic stimulus, pro-apoptotic proteins are released and free to induce MOM permeabilization. Interactions among Bcl2 family members that control the fate of the mitochondria have been thought to occur through soluble domains, especially BH domains(15). However, recent findings suggest that their TMDs also participate in these PPIs(16–18) and that these intramembrane interactions are crucial for apoptotic control(18).

Here, we developed a novel rational methodology to design sequences that specifically recognize the TMD of natural proteins. To illustrate this approach we explored the intramembrane protein-protein interactions (PPIs) of BclxL, an anti-apoptotic member of the Bcl2 protein family. Next, we designed sequences capable of selectively sequestering BclxL TMD and limiting the anti-apoptotic effect of BclxL. We hope that this work sparks the development of a new generation of inhibitors targeting interactions among transmembrane domains.

## Results

### The TMD of BclxL establishes interactions with pro- and anti-apoptotic Bcl2 members

Bcl2 and Mcl-1 can establish intramembrane interactions with other members of the Bcl2 family(16, 17). Furthermore, to block apoptosis, viral analogs of Bcl2 proteins must interact through the TMD with cellular Bcl2 members(18). We sought to explore whether the BclxL TMD can be used to establish interactions with other pro- and anti-apoptotic Bcl2 members. To assess these potential intramembrane contacts, we employed BLaTM, a genetic tool designed to qualitatively and quantitatively study TMD– TMD interactions in *E. coli*(*19*). Briefly, the tested TMDs are fused to either the Nt or the Ct end of a split β-lactamase (βN and βC, respectively) and the enhanced green fluorescent protein (eGFP) (Figure 1a). Additionally, the chimeras include the pelB cleavable signal peptide, which directs the protein to the inner bacterial membrane and determines its topology, ensuring periplasmic localization of the β-lactamase. In the inner membrane, an efficient TMD–TMD interaction facilitates the reconstitution of the β-lactamase and thus the growth of bacteria in selective media (ampicillin) (Figure 1a). In this assay, the LD_50_ of the antibiotic serves as an indicator of the strength of the assayed TMD–TMD interaction, while the eGFP-derived fluorescence allows for rapid quantification of protein levels. As a positive control and for normalization purposes, we used the TMD of glycophorin A (GpA), a hydrophobic segment that can form noncovalent homodimers within the membrane(20–22). The non-oligomerizing TMD of the mitochondrial protein Tomm20 (T20) was used as a negative control for membrane overcrowding and stochastic interactions(18). Of note, bacteria can grow only in the presence of ampicillin when the β-lactamase is reconstituted in the periplasm, i.e., only if the tested regions are properly inserted in the bacterial inner membrane. Therefore, the BLaTM assay also indicates the insertion potential of the tested regions. Using this approach, we tested the homo-oligomerization of BclxL TMD and its hetero-oligomerization with the TMD of anti-apoptotic Bcl2, and the TMDs of pro-apoptotic (Bax and Bak) and BH3-only (Bik and Bmf) Bcl2 members. Our results indicated that BclxL TMD forms a weak homo-oligomer (Figures 1b and 1c). Furthermore, we identified transmembrane hetero-oligomers with Bcl2, Bax, and Bak.

**Figure 1.**
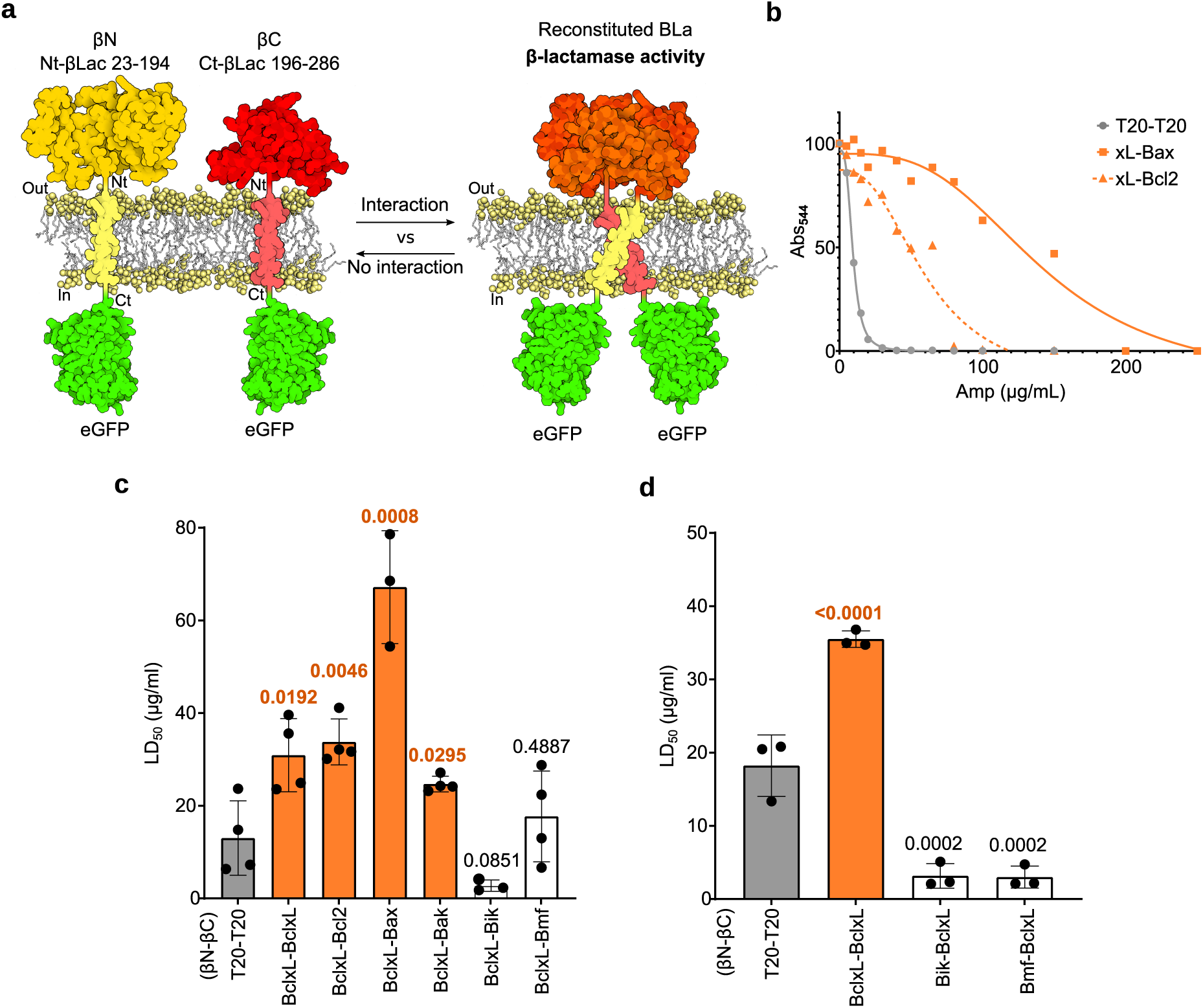
BlcxL’s TMD interactions in biological membranes. **a**. Schematic representation of the BLaTM assay. The β-Lactamase was split in two non-active fragments (Nt-βLac and Ct-βLac). Each of them was fused to a TMD and the eGFP in C-terminus generating the βN and βC chimeras respectively. A TMD–TMD interaction approximates the Nt-βLac and Ct-βLac fragments and facilitates the reconstitution of the β-Lactamase structure and activity (ampicillin resistance). The residues constituting the Nt-βLac and Ct-βLac halves are specified above the protein representation. **b**. In the BLaTM assay, the strength of a TMD–TMD interaction correlates with resistance to ampicillin. The panel shows representative examples of the dose-response curves from which the ampicillin LD_50_ was calculated. The dose-response curves corresponding with the interactions between the TMDs of BclxL and Bax (orange, squares), BclxL and Bcl2 (orange, triangles), and the TMD of T20 are shown. **c**. and **d**. The βN and βC chimeras bearing the TMD of the indicated proteins were co-expressed in E. coli and the resulting ampicillin LD_50_ was measured. The βN T20-βC T20 homodimer was used as a negative control (gray), and the βN GpA-βC GpA homodimer was used as a positive control and normalization value across experimental replicates (LD_50_∼100 μg/mL). The normalized means and standard deviations of at least three independent experiments (n ≥ 3) are shown. The individual value for each experiment is represented by a solid dot. An interaction (highlighted in orange) was considered if the observed LD_50_ was significantly higher (two-tailed homoscedastic t-test, p-value < 0.05) than the negative control (gray bar). P-values are indicated above the corresponding bar.

The reporter signal resulting from a TMD–TMD interaction depends not only on the sequences of the interacting TMDs and thus their inherent affinity, but also on the orientation of the interacting surfaces of the TMDs in relation to their accompanying signaling domains(18, 19). To confirm that the lack of interaction between the TMD of BclxL and the TMDs of Bik and Bmf was not the result of misaligned reporter domains(19) we tested opposing combinations of BLaTM chimeras, i.e., βN-Bik/βC-BclxL and βN-Bmf/βC-BclxL (Figure 1d). Despite the change in the experimental set-up, we observed no interaction between the BclxL TMD and the TMDs of Bik or Bmf. Of note, expression levels, measured using GFP fluorescence, were comparable for all BLaTM chimeras (Supp. Figure 1). Next, to corroborate the interaction capabilities of the BclxL TMD in eukaryotic cells, we used a bimolecular fluorescent complementation (BiFC) aproach(23), adapted for the study of intramembrane interactions(16, 18, 24). Briefly, the tested TMDs were fused to a split Venus fluorescent protein (VFP), either to its N-terminus (VN) or its C-terminus (VC), neither of which is fluorescent. Interaction of the TMDs brought together the VN and VC ends, reconstituting the VFP structure and restoring its fluorescence (Figure 2a). As noted, the TMD of GpA was used as a positive control and normalization value across experimental replicates, and the TMD of T20 was used as a negative control. Our results indicated that the BclxL TMD can homo-oligomerize in eukaryotic membranes. Furthermore, using BiFC, we detected hetero-oligomers with the TMDs of Bcl2, Bak, and Bik (Figure 2b). We also tested the VN Bax-VC BclxL and VN Bmf-VC BclxL combinations of BiFC chimeras (Figure 2c). Despite the change in the experimental set-up, no interaction was observed between the TMD of BclxL and the TMDs of Bax or Bmf. Of note, western blot analysis indicated that the BiFC chimera bearing the Bax TMD was consistently expressed at lower levels, which could explain the differences seen with the BlaTM assay (Figure 2d). For a more comprehensive visualization of the aforementioned interactions, we also have included a network representation (Supp. Figure 2).

**Figure 2.**
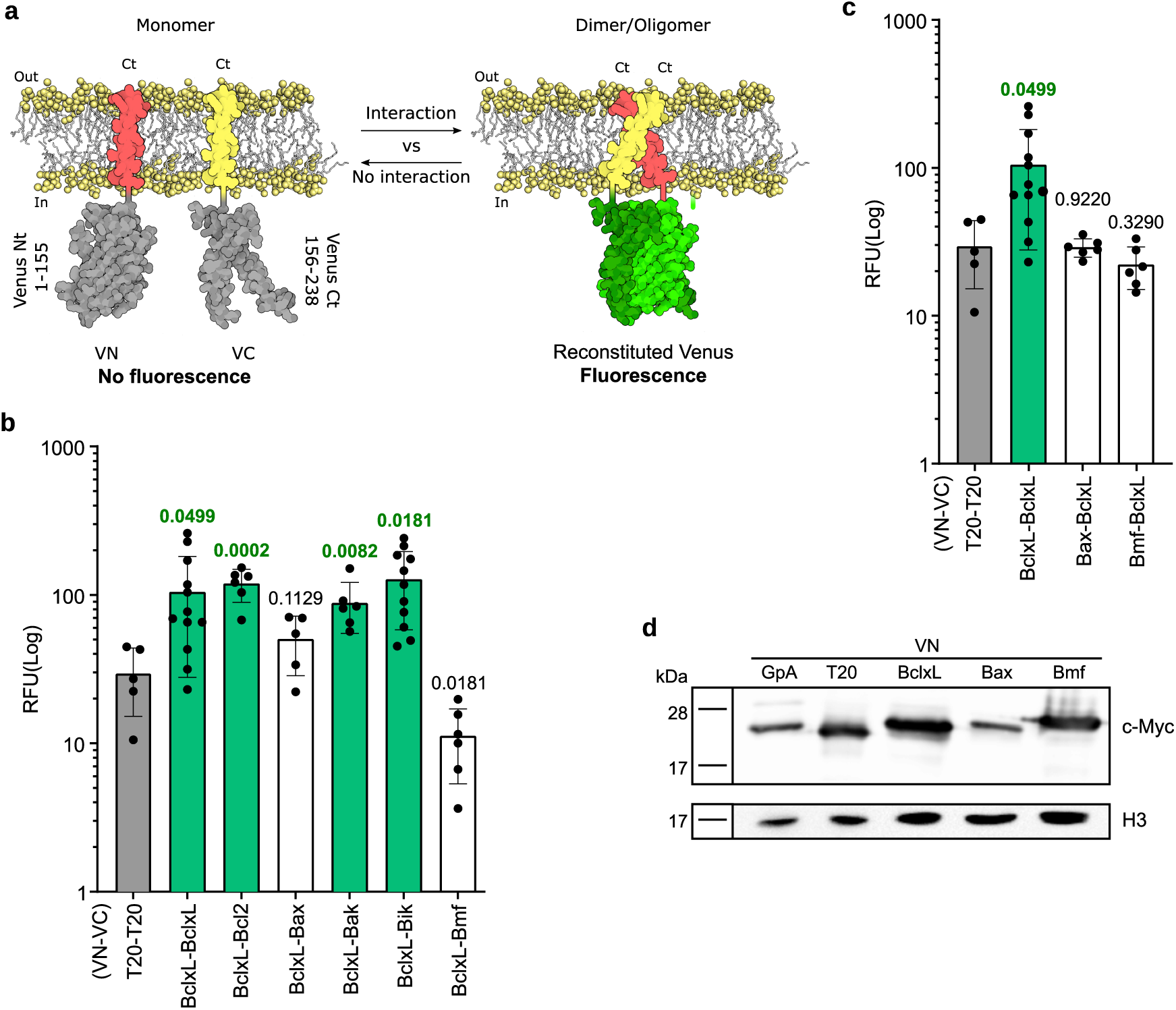
Interactions of the BclxL TMD in eukaryotic membranes. **a**. Schematic representation of the Bimolecular Fluorescent Complementation (BiFC) assay. The two non-fluorescent fragments of a split Venus fluorescent protein (Venus Nt and Venus Ct; both in gray) are fused to two potential interacting TMDs (yellow and red) rendering two chimeric constructs designated respectively as VN and VC. The association of the TMDs facilitates the reconstitution of the fluorescent protein structure (green). The residues included in each VFP fragment are indicated next to the protein representation. **B** and **c** Relative fluorescence units (RFU) for the homo-oligomerization of the TMD of BclxL and its hetero-oligomerization with the TMD of Bcl2, Bax, Bak, Bik, and Bmf; the mean and standard deviation (n ≥ 5) are shown. Solid dots represent the results of individual experiments. The TMD included in each chimera (either VN or VC) is indicated below each bar. The VN T20-VC T20 combination was used as negative control (gray bar). The VN GpA-VC GpA was used as a positive control and normalization value across experiments. An interaction (highlighted in green) was considered to have occurred only if the obtained RFU was significantly higher (two-tailed homoscedastic t-test, p-value < 0.05) than the negative control, p-values are indicated above the corresponding bars. **d**. Western blot analysis of VN-GpA, VN-T20, VN-BclxL, VN-Bax, and VN-Bmf protein levels. Histone 3 (H3) was used as a loading control. Experiments were done in triplicates.

### BclxL intramembrane interactions are crucial for its anti-apoptotic role

Next, to investigate whether these newly found TMD–TMD interactions are necessary for BclxL control of cellular apoptosis, we transfected HeLa cells with BclxL with or without the TMD (BclxL-FL and BclxL-ΔTMD, respectively). Additionally, we included a chimera in which the TMD of BclxL was replaced by the TMD of T20 (BclxL-T20). All of these constructs included a Nt c-myc tag to facilitate detection. As a negative control, cells were transfected with an empty plasmid (Empty). Once transfected, cells were treated with doxorubicin to induce apoptosis(18, 25) and the percentage of surviving cells 16 h post-treatment was calculated based on Trypan blue staining (Figure 3a and b). Our results indicated that when doxorubicin is used as a cell death stimulus, BclxL requires the TMD to block apoptosis. Furthermore, the substitution of this TMD by the TMD of T20 impaired the anti-apoptotic function of BclxL, suggesting that intramembrane interactions of BclxL with other Bcl2 members are necessary for BclxL’s anti-apoptotic function. Western blot analysis confirmed comparable expression levels for the BclxL-FL, BclxL-ΔTMD, and BclxL-T20 variants (Figure 3c).

**Figure 3.**
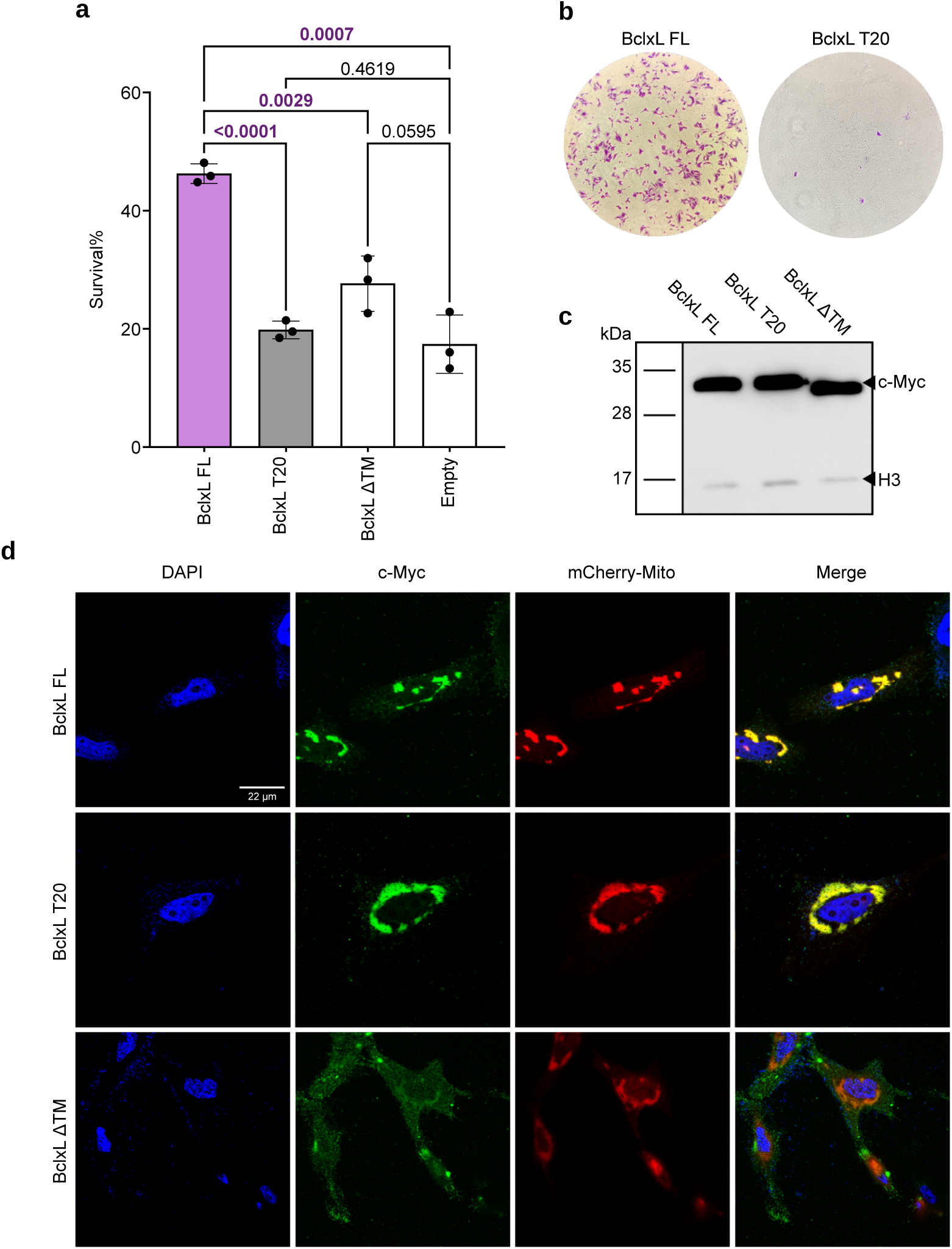
The importance of the BclxL TMD in apoptotic control. **a**. Survival of doxorubicin-treated cells. HeLa cells were transfected with BclxL with or without the TMD (BclxL-FL and BclxL-ΔTMD, respectively), we also included a chimera in which the TMD of BclxL was replaced by the TMD of T20 (BclxL-T20). Next, cells were treated with doxorubicin [15 µM]final, and the percentage of surviving cells 16 h post-treatment was calculated based on Trypan blue staining. The survival percentage means and standard deviations of three independent experiments are shown (n = 3). Solid dots represent the results of individual experiments. Transfection with an empty plasmid (Empty) was used as a negative control. The level of significance (ordinary one-way ANOVA test with Dunnett correction, p-value < 0.05) is shown, and significant differences are highlighted in green bold letters. **b.** Alternatively, cells were transfected with Bclxl FL or BclxL T20 and treated with doxorubicin. Approximately 16 h post-treatment cells were washed, fixed, and stained. Representative images are included. Images were taken using a 10x microscope objective. **c**. Western blot analysis of BclxL-FL, BclxL-ΔTM, and BclxL-T20 protein levels. Histone 3 (H3) was used as a loading control. A representative assay (n = 3) is shown. **d**. Subcellular localization of BclxL chimeras. BclxL-FL, BclxL-ΔTM, and BclxL-T20 with a c-Myc tag were transfected into HeLa cells together with a mitochondrial marker (mCherry-mito, red). After 24 h, cells were immunostained using an anti-c-Myc antibody (green). DAPI staining is shown in blue. The right column of each panel shows the co-localization of the BclxL chimeras and the mitochondrial marker (yellow), visible only when the images are merged. Experimental n = 3.

BclxL localizes primarily in the mitochondria(26–28). Given the importance of TMDs for membrane protein sorting(29), particularly in the case of tail-anchoring proteins(30), we thought it was important to consider how deletions or substitutions in the Ct hydrophobic region of BclxL would affect its cellular localization. To analyze the subcellular location of the previously mentioned chimeras, BclxL (BclxL-FL) and the BclxL-T20 and BclxL-ΔTMD variants were expressed in HeLa cells alongside a fluorescent mitochondrial marker (mCherry-Mito, Figure 3d). The fluorescence micrographs revealed a similar subcellular distribution between BclxL-FL and BclxL-T20. Furthermore, both moieties were located at the mitochondria, as indicated by co-localization with the mitochondrial marker. BclxL-ΔTMD, however, showed a different cellular distribution. Removing the TMD left the protein in the cytosol, impeding its co-localization with the mitochondrial marker.

### Computational design of BclxL inhibitors

Once we had established the importance of BclxL TMD–TMD interactions for the anti-apoptotic function, we sought to design an inhibitor that could block these intramembrane PPIs. Such an inhibitor should, in principle, block the pro-survival effect of BclxL and thus sensitize cells to an apoptotic stimulus.

Inhibitor design started with the modeling of the BclxL homo-interaction using TMHOP (Trans-membrane Homo Oligomer Predictor)(5). TMHOP uses Rosetta symmetric all-atom ab initio fold-and-dock simulations in an implicit membrane environment to predict thousands of low-energy conformations based on the energy function described by Weinstein et al.(5). This energy function relies on the empirical measurement of amino acid insertion propensities into the bacterial inner membrane(6). It encodes a lipophilicity term for each amino acid, depending on each amino acid’s exposure to the lipid and its distance from the cytosolic or extracellular milieu. TMHOP generates several models for the desired interaction (Figure 4a). Based on structural characteristics and associated Rosetta energy, we selected a TMHOP model that forms a tightly packed parallel dimer. The structure of the dimer and the monomer that constitutes it can be found in Figure 4b. Roughly half of each monomer’s surface area (2389.5Å^2^) is in contact with its companion (1244.1Å^2^). The amino acids involved in the contact between monomers are shown in Figure 4 c. The crossing angle of the dimer was exactly 48°, with the shortest distance (4.1Å) between the N atoms of Met at position 9. To avoid confusion, we numbered the amino acids based on their position within the TMD.

**Figure 4.**
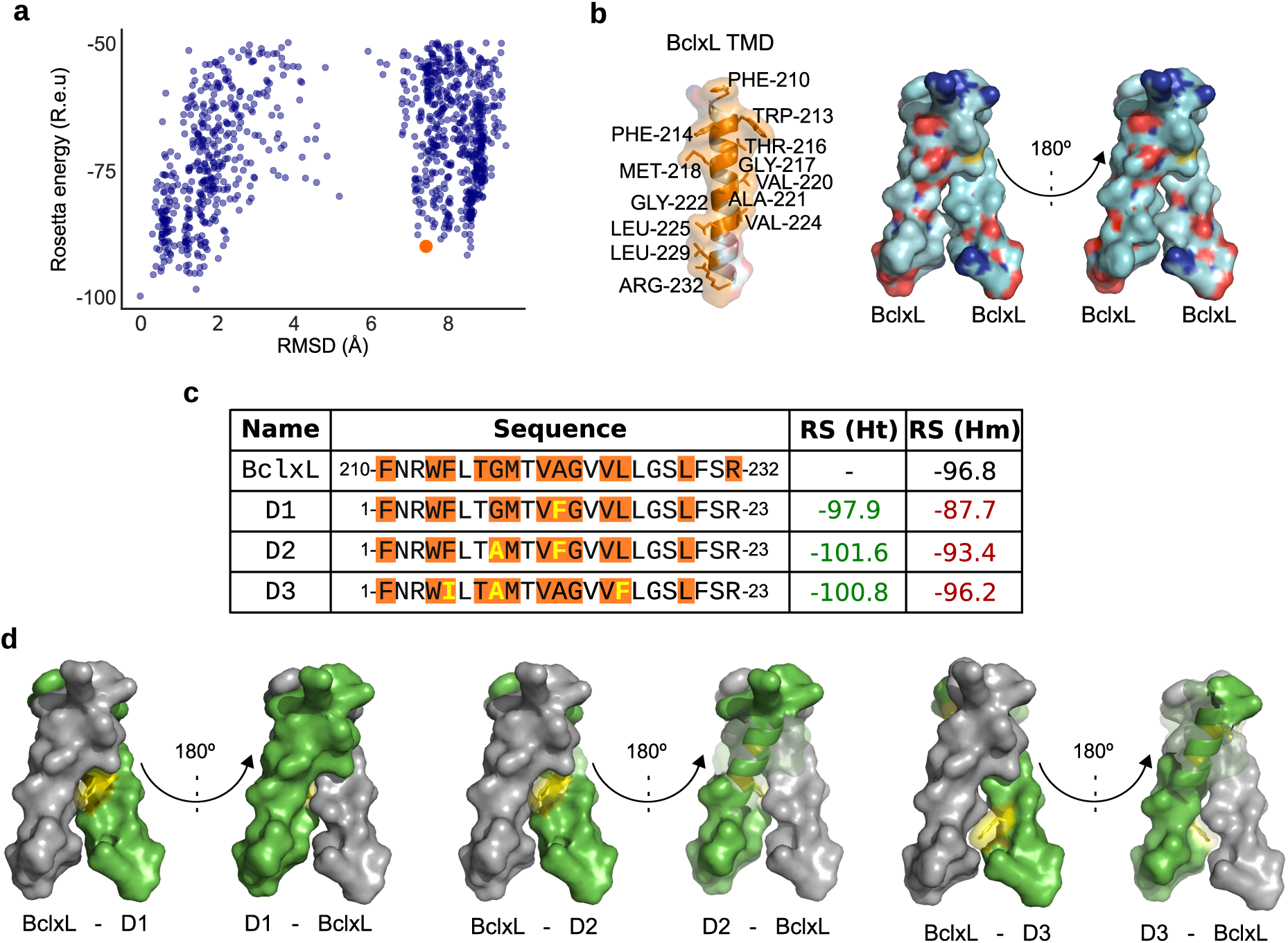
Design of BclxL TMD inhibitors. **a**. Scatter plot distribution for the BclxL TMD homodimer modeling. Dots represent each of the models for the BclxL TMD’s homodimers. RMSD is calculated from the lowest energy model. The selected model is highlighted in orange. b. Structural representations of the selected model. The structure of the monomer and the dimer (frontal and 180° turn views) are shown. The monomer representation features the amino acids on the interacting surface, numbering based on the BclxL sequence. In the dimer representation, the nitrogen and oxygen atoms are highlighted in blue and red respectively c. Sequences of the TMDs of the BclxL and the D1, D2, and D3 designs. Differences among the sequences are featured in yellow. The amino acids in the interaction surfaces of the potential dimers between the TMD of BclxL (gray) and the D1, D2, and D3 designs (green) are shown in orange. The Rosetta scores for the hetero-(RS Ht) and homodimerizations (RS Hm) are indicated. Values lower and higher than RS(Hm) for BclxL TMD are shown in green and red respectively. The positions of the amino acids in the TMD of BclxL (210–232) and the designs (1–23) are included. d. Frontal view (and 180° turn) of the potential dimer between the TMD of BclxL (gray) and the D1, D2, and D3 designs (green). Changes in the designs over the TMD sequence of BclxL are highlighted in yellow.

The selected model was fed into the FuncLib design algorithm(31) to generate higher affinity binders that could serve as inhibitors. The FuncLib algorithm uses Rosetta design calculations to enumerate combinations of tolerated amino acid substitutions at specific positions. It then relaxes each combination using whole-protein minimization (based on the Rossetta membrane energy function(5)) and ranks these combinations by energy. In addition to the atomistic design steps, FuncLib uses information from multiple-sequence alignments to eliminate mutations that are not commonly observed in homologs. This method has been successfully applied to dramatically improve enzyme catalytic rates(31), binding affinity(32), and antibody stability and affinity(33), but it has not been applied previously to membrane proteins. Additionally, single-span TMDs tend to self-associate(24), whereas our design objective was an inhibitor that would target BclxL specifically and exhibit minimal self-association propensities. To do so we computationally modeled each design to obey the following rules. Positive selection of a heterodimer (Non-symmetric FuncLib; ΔΔG<+1 Rosetta energy units; R.e.u.); Negative selection for a homodimer (symmetric FuncLib; ΔΔG>+5 R.e.u.).

The resulting potential TMD inhibitors were manually curated after visualization of the 3D structural models, and we selected three designs (designated as D1, D2, and D3). The sequences of each selected model, as well as the BclxL TMD, together with their corresponding Rosetta energy for their homo- and hetero-dimerization, are shown in (Figure 4c). Structural representations of the interactions among the designs (D1, D2, and D3) and the TMD of BclxL are given in Figure 4d. Additionally, the amino acids found in the contact surface, the contact area between monomers, and their crossing angles are shown in Supp. Figure 3.

### Membrane insertion potential of D1, D2, and D3

Before testing the inhibition properties of the designed sequences, we verified that these segments could be inserted into the membrane together with BclxL. To that end, we tested the membrane insertion of the D1, D2, and D3 sequences into ER-derived microsomes using an *in vitro* transcription/translation assay. This assay, designed for an accurate and quantitative description of the membrane insertion capability of short sequences, is based on the *E. coli* leader peptidase (Lep). Lep is an Nt/Ct luminal membrane protein consisting of two TMDs (H1 and H2) connected by a cytoplasmic loop (P1) and a large C-terminal domain (P2) (Supp. Figure 4a). We used a Lep-derived system (Lep′) containing an extended Nt(34–36). Additionally, the Lep’ includes two N-glycosylation acceptor sites consisting of an Asn-X-Ser/Thr sequence, where X can be any amino acid except Pro(37). The first acceptor site (G1) is in the extended Nt, whereas the second site (G2) is in the P2 domain. N-linked glycosylation has been extensively used as a topological reporter for more than two decades(38). This post-translational modification occurs only in the ER lumen, the location of the active site of oligosaccharide transferase, a translocon-associated enzyme responsible for oligosaccharide transfer(39); thus, no N-linked glycosylation occurs in polypeptide sequences spanning the membrane or facing the cytosol. In Lep’, the hydrophobic region to be tested replaced the H2 domain (Supp. Figure 4a). When the tested hydrophobic region is recognized by the translocon as a TMD and inserted into the membrane, both G1 and G2 are oriented towards the ER lumen, yielding a doubly-glycosylated version of Lep’. If the tested region is not inserted into the membrane, however, G2 stays on the cytoplasmic side and is not modified by the oligosaccharide transferase. Glycosylation of an acceptor site increases the molecular mass of the protein by ≈ 2.5 kDa relative to the observed molecular mass in the absence of membranes, allowing monitoring of the glycosylation state (one vs. two glycosylations) and thus insertion into the membrane of the tested sequence (Supp. Figure 4a right).

According to our results, all three designed sequences (D1, D2, and D3) are efficiently inserted into the membrane, as indicated by the double glycosylation pattern (Supp. Figure 4b). In this assay, for control sequences, we used the hydrophobic regions of the Bcl2 proteins Mcl1 (insertion control) and Noxa (no-insertion control)(17, 40). The experimentally obtained ΔGs (ΔG_app_^exp^) correlated with the *in silico* insertion potential of the designed sequences (Supp. Figure 4b), based on values calculated using the *ΔG prediction server*(*41, 42*) with the default parameters. Of note, the ΔG_app_^pred^ for any of the three designs was lower than the ΔG_app_^pred^ associated with the TMD of BclxL (−0.355 kcal/mol).

### Analysis of the interaction between D1, D2, or D3 with the TMD of BclxL

Next, using BLaTM, we analyzed the interactions between the TMD of BclxL and the computationally designed inhibitors D1, D2, and D3 (Figures 5a, 5b). The results of these experiments revealed that D1 can efficiently bind to the BclxL TMD but does not form homo-oligomers, as we intended in our design. Of note, the interaction between the TMD of BclxL and D1 was stronger than the homo-oligomerization of the TMD of BclxL. Although D2 and D3 did not form homo-oligomers, they did not interact with the TMD of BclxL (Figure 5b), regardless of the assayed combination (Figure 5c). Thus, we found no significant differences among the average LD_50_ values associated with the BclxL TMD-D2, BclxL TMD-D3, and T20-T20 BLaTM assays. Expression levels (measured using eGFP fluorescence) were comparable for all βN and βC chimeras (Supp. Figure 5).

**Figure 5.**
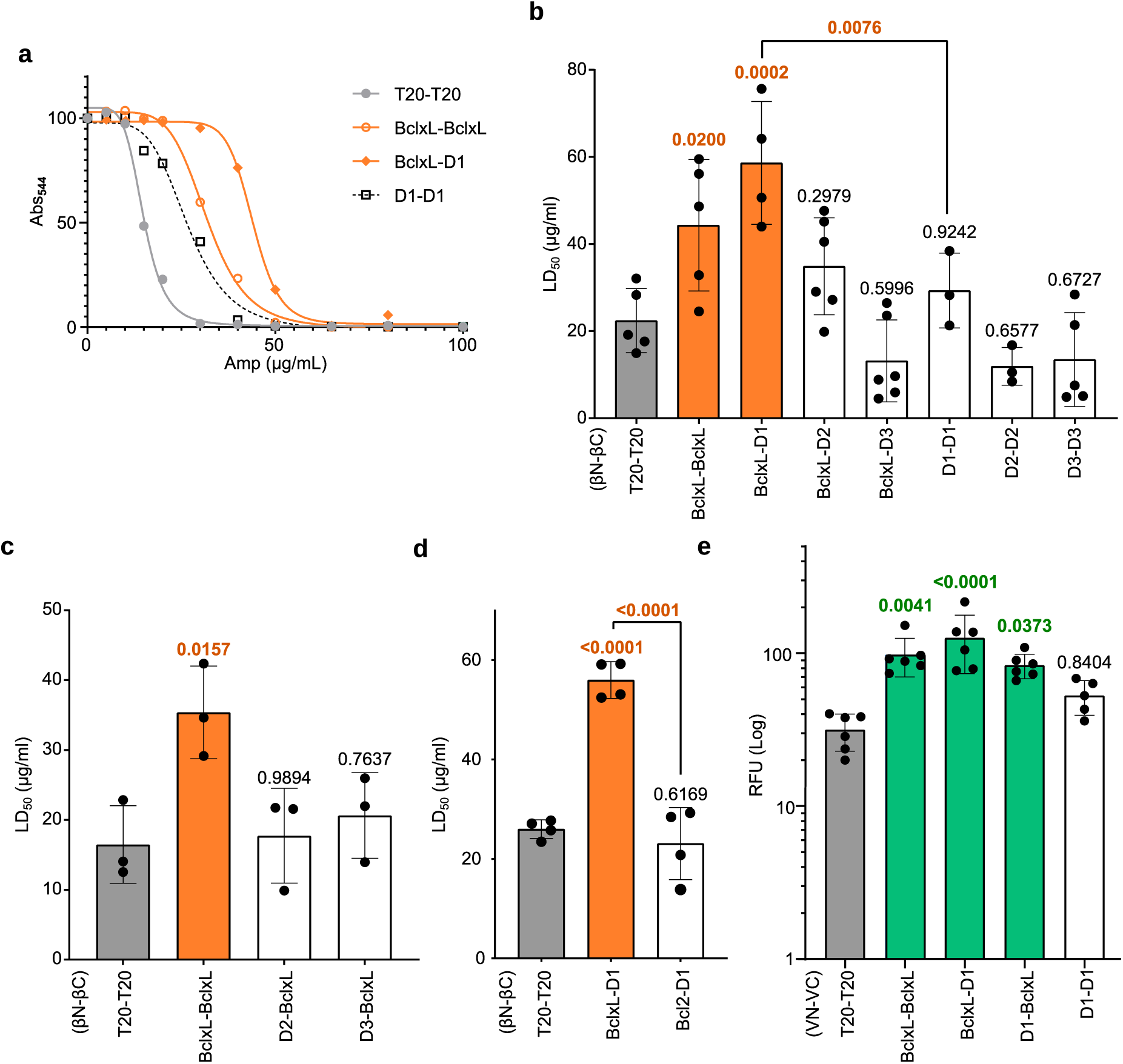
Interaction between the TMD of BclxL and the designed inhibitors. **a**. Representative example of ampicillin dose-response curves in the BLaTM assay. The TMD of T20 was used as negative control (gray). The TMD of GpA was used as positive control and normalization value across experiments. The BclxL-BclxL (orange circles), BclxL-D1 (orange diamonds), and D1-D1 (black dashed line, squares) interaction profiles are shown. **b**, **c**, and **d**. Interaction between the TMD of BclxL and the designed inhibitors. BlaTM chimeras (βN-βC) bearing the indicated TMD were co-expressed in *E. coli* and the resulting ampicillin LD_50_ was measured. The means and standard deviations of at least three independent experiments are shown. Individual values for each experiment are represented by solid dots. The βN T20-βC T20 homodimer was used as a negative control (gray). The βN GpA-βC-GpA homodimer was used as a positive control (not included). An interaction (highlighted in orange) was considered if the obtained LD_50_ was significantly higher (ordinary one-way ANOVA test with Dunnett correction, p-value < 0.05) than the negative control. P-values are indicated above the corresponding bar. The level of significance (ordinary one-way ANOVA test with Dunnett correction, p-value <0.05) when comparing xL-D1 vs D1-D1 is included. **e**. BiFC analysis of the interaction between the TMD of BclxL and D1 in eukaryotic membranes. The TMD included in each chimera (VN or VC) is indicated. The RFU mean and standard deviation of at least five independent experiments are shown. Solid dots represent the results of individual experiments. The TMD of T20 was used as negative control (gray). The TMD of GpA was used as a positive control and normalization value across experiments. An interaction (highlighted in green) was considered if the obtained RFU was significantly higher (two-tailed homoscedastic t-test, p-value < 0.05) than the negative control (gray bar). P-values are indicated above the corresponding bar (significantly higher in bold green).

Next, we investigated the specificity of the observed interaction by challenging D1 with the TMD of Bcl2, another anti-apoptotic protein. To do so, we once again used the BLaTM approach and tested the BclxL TMD-D1 and Bcl2 TMD-D1 interactions side by side. We detected no interaction between D1 and the Bcl2 TMD (Figure 5d). Thus, any effect of D1 on cell survival would most likely arise from its interaction with the BclxL TMD. The similar expression levels of all BLaTM chimeras used in this assay (Supp. Figure 5) indicated that the observed differences were not the result of different protein concentrations within the cells.

Additionally, we used the BiFC assay to ensure that the interaction between D1 and the BclxL TMD was maintained in eukaryotic membranes (Figure 5e). The results indicated that D1 could efficiently bind to the TMD of BclxL but did not form homo-oligomers. Of note, the D1 and BclxL TMD hetero-interaction was identified regardless of the combination of BiFC chimeras analyzed.

D1 and D2 share a single amino acid change over the TMD sequence of BclxL, specifically Ala12Phe (Figure 4c). Of note, this is the only difference between D1 and the TMD of BclxL. In light of these data, we decided to test whether other amino acid substitutions in the same position would facilitate the interaction with the BclxL TMD while also avoiding homo-interactions. First, we analyzed which substitutions at position 12 of the TMD of BclxL would allow the insertion of the resulting sequence into the mitochondrial membrane. For this purpose, we used the ΔG prediction server(41, 42) (Supp. Figure 6a). The amino acid substitutions that kept the ΔGpred of the resulting segment below zero (indicative of membrane insertion) were included in the BLaTM assay for assessment of their homo-oligomerization and hetero-oligomerization (with the TMDs of BclxL and Bcl2) properties. Only one of the tested substitutions (Ala12Cys) facilitated the interaction with the BclxL TMD while avoiding the formation of homo-oligomers or off-target PPIs with the TMD of Bcl2 (Supp. Figure 6b and c). However, in eukaryotic membranes, the Ala12Cys substitution did not preclude the formation of homo-oligomers (Supp. Figure 6d) ruling out its use as a BclxL inhibitor.

### Subcellular localization of BclxL inhibitors

To inhibit the anti-apoptotic effect of BclxL, the designed sequences must be located in the same cellular compartment where BclxL is found. To test their location, we fused D1 and D2 sequences to the Ct of the eGFP (eGFP-D1 and eGFP-D2) and expressed this construct in HeLa cells together with BclxL attached to the fluorescent protein mCherry (mCherry-BclxL). D3 was excluded from the assay because of the poor interaction with BclxL TMD observed in the BLaTM assay. We intentionally fused the designed sequences at the Ct of the eGFP to mimic the topology of a protein of the Bcl2 family and consequently its localization(43). Next, we analyzed the subcellular distribution of these chimeras by confocal fluorescence microscopy (Figure 6a). Additionally, chimeras bearing the TMD of BclxL or T20 were included in the assay as controls (eGFP-xL and eGFP-T20, respectively). As expected, eGFP-xL and mCherry-BclxL had the same subcellular distribution. mCherry-BclxL and eGFP-T20 were also co-localized in HeLa cells. Analysis of the locations of eGFP-D1 and eGFP-D2 showed a strong co-localization with mCherry-BclxL (Figure 6), indicating a similar cellular distribution for the D1 and D2 inhibitors and BclxL. We did not observe co-localization of the mCherry-BclxL with eGFP bearing no TMD (Supp. Figure 7).

**Figure 6.**
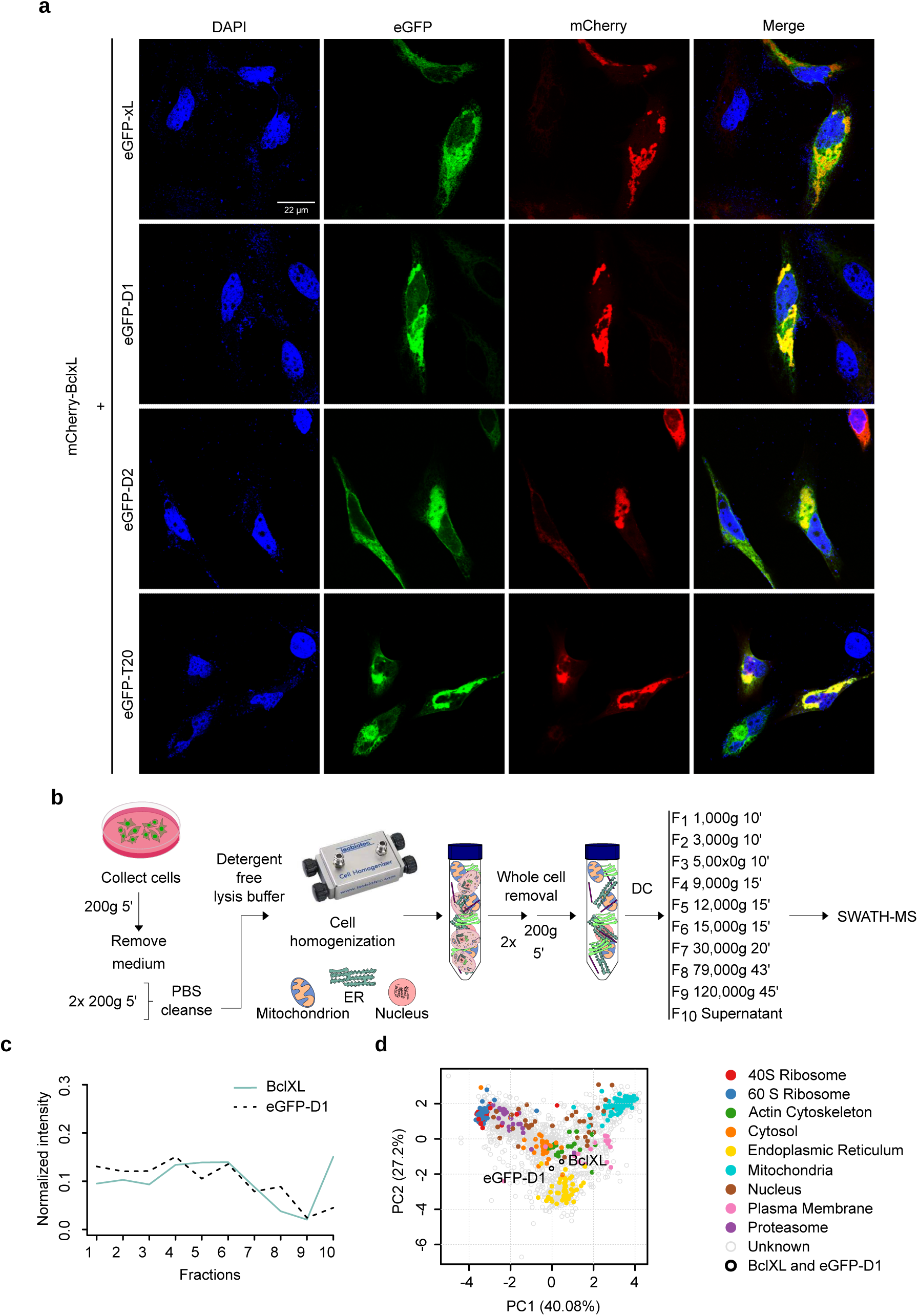
Subcellular localization of the designed inhibitors. **a**. The eGFP was fused to the TMD of BclxL, D1, D2, or the TMD of T20 to create the eGFP-xL, eGFP-D1, eGFP-D2, and eGFP-T20 chimeras, respectively. These chimeras (green) together with a construct bearing BclxL fuse to the mCherry fluorescent protein (mCherry-BclxL, red) were transfected into HeLa and 24 hours later analyzed by confocal microscopy (n=3). DAPI staining is shown in blue. The right column of each panel shows the co-localization of both signals (yellow, visible only when the images are merged). **b**. Overview of the organelle differential centrifugation workflow. In the assay, a series of differential ultracentrifugation steps was used to separate organelles and subcellular compartments. The proteins in each fraction were quantified by SWATH-MS analysis. **c**. Abundance profiles of BclxL (blue line) and eGFP-D1 (black dotted line) across all fractions. A representative result is shown. **d**. Principal Component Analysis of the abundance profiles across fractions obtained by differential centrifugation. The plot shows the 1962 proteins identified by SWATH-MS. The percentage in each axis represents the amount of total variability that PC1 and PC2 can explain. Organelle markers are colored. eGFP-D1 and BclxL are shown in white with a black contour, and the rest of the proteins (other) are shown in gray. A representative plot is shown.

To ensure that D1 and BclxL coexist in the same cellular compartment and thus that D1 could block the anti-apoptotic role of BclxL, we performed a localization assay based on organelle differential ultracentrifugation(44). Briefly, cells expressing either BclxL or eGFP-D1 were homogenized in a detergent-free medium to avoid organelle content release. Next, the cell lysate was divided into 10 fractions by sequential differential centrifugation, as previously described(44) (Figure 6b). We then analyzed the protein content in each fraction by sequential window acquisition of all theoretical mass spectra (SWATH-MS)(45, 46). Data analysis and visualization, as well as protein localization, were done with a dedicated open-source R package(47, 48). BclxL and eGFP-D1 had a similar distribution profile, suggesting a similar subcellular localization (Figure 6c and 6d, Supp. Figure 8). Minor differences were found at fraction 10, suggesting that Bclx exists partially as a cytosolic protein(49–51). A comparison of D1 and BclxL profiles with a set of organelles markers revealed no distinct protein localization for either of these proteins (Supp. Figure 8 and 9). Of note, similar results have been obtained previously for Bcl2(52), Bax(44), and mouse BclxL(53). A list of the organelle markers together with the SAWTH-MS data can be found in the supplementary information.

### D1 and D2 inhibit the anti-apoptotic effect of BclxL

Finally, we tested the anti-apoptotic effect of D1 and D2. HeLa cells were transfected with BclxL alongside the eGFP-T20, eGFP-D1, eGFP-D2, or eGFP-xL chimeras. As a control, we used cells that did not receive BclxL or any of the chimeras and transfected them with an empty plasmid (Empty) to keep the amount of transfected DNA constant across all samples. After transfection, cells were treated with doxorubicin to induce apoptosis. The cells that received eGFP-T20 or eGFP-xL plus BclxL could block doxorubicin-induced apoptosis (Figures 7a and b). Remarkably, transfection of eGFP-D1 eliminated the anti-apoptotic effect of BclxL. D2 also reduced cell viability but less drastically than D1. Of note, no significant differences were found between the samples transfected with eGFP-D1 or eGFP-D2 or cells transfected with an empty plasmid when treated with doxorubicin, indicating that both D1 and D2 are capable of inhibiting BclxL function (Figure 7a). Western blot analysis confirmed comparable expression levels for the eGFP-T20, eGFP-D1, eGFP-D2, and eGFP-xL chimeras (Figure 7c). Furthermore, the fluorescence levels of all four eGFP chimeras were comparable (Figure 7d).

**Figure 7.**
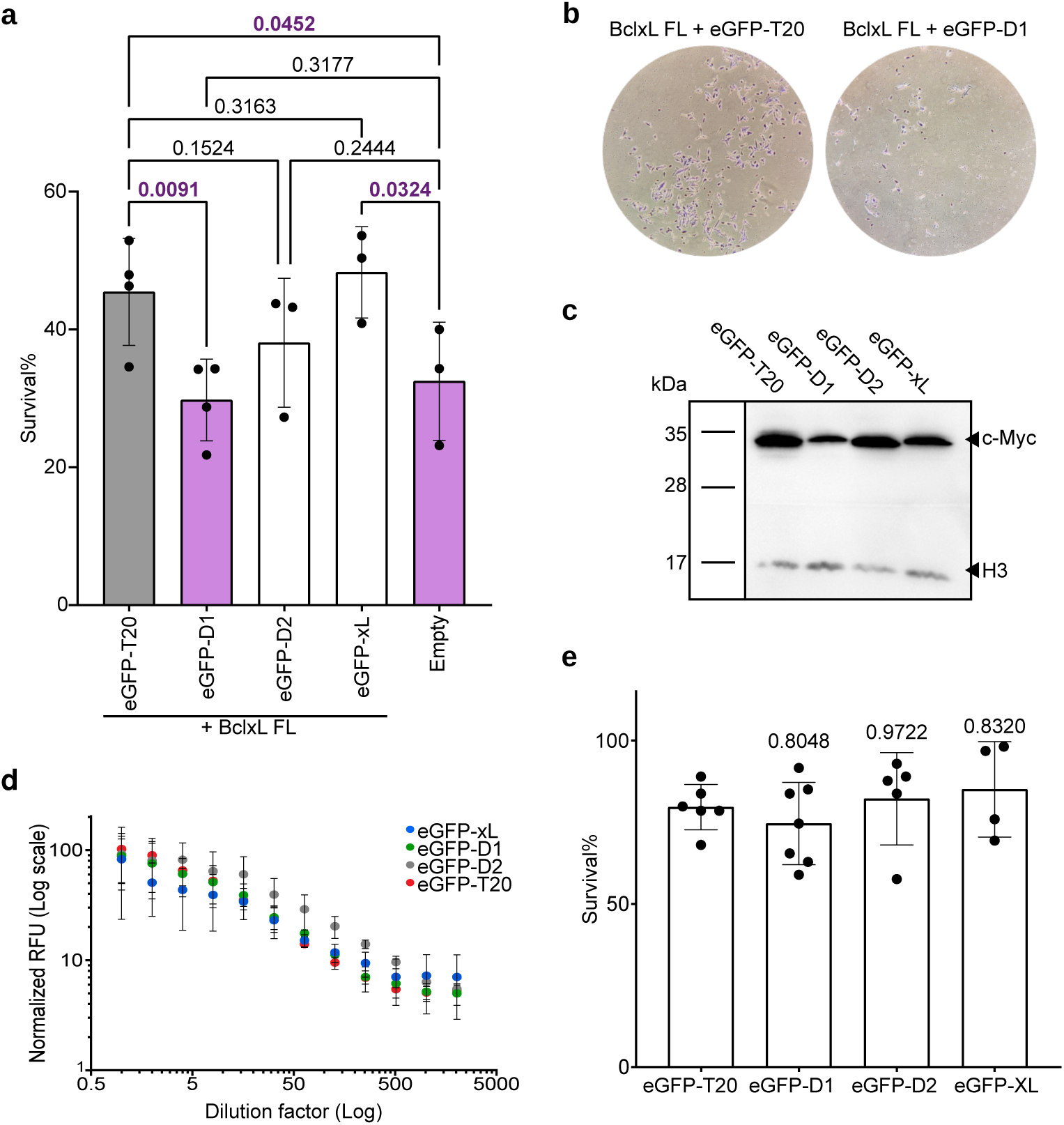
Inhibition of the BclxL anti-apoptotic effect. **a**. HeLa cells were transfected with BclxL together with either the eGFP-XL, eGFP-D1, eGFP-D2, or eGFP-T20 (gray bar, negative control) chimeras. Next, cells were treated with doxorubicin, and after 16 h the percentage of surviving cells was calculated based on Trypan blue staining. The survival percentage mean and standard deviation of at least three independent experiments are shown. Solid dots represent the results of individual experiments. Transfection with an empty plasmid (Empty; yellow) was used as a negative control. Statistical differences are based on a one-tailed homoscedastic t-test (p-values are indicated above). Values significantly lower than those obtained with the T20 control are highlighted in green. **b.** HeLa cells were co-transfected with Bclxl FL + eGFP-D1 or BclxL Fl + eGFP-D20 and treated with doxorubicin. After 16 h cells were washed, fixed, and stained. Images were taken using a 10x microscope objective. **c**. Western blot analysis of protein levels. Histone 3 (H3) was used as a loading control (n = 3). **d**. Fluorescence levels of the eGFP chimeras. HeLa cells were transfected with the indicated eGFP chimeras. After 24 hours samples were serially diluted and the relative fluorescence units (RFUs) and measured (λexc 485 nm, λem 535nm). The mean and standard deviation of at least three independent experiments are shown. **e**. HeLa cells were transfected for analysis of the toxicity associated with the expression of the eGFP-XL, eGFP-D1, eGFP-D2, or eGFP-T20 chimeras. The percentage of surviving cells was calculated based on Trypan blue staining 24 hours after transfection. The survival percentage mean and standard deviation of at least five independent experiments are shown. Solid dots represent the results of individual experiments. For statistical purposes, eGFP-T20 was used as a control. Statistical differences are based on a two-tailed homoscedastic t-test, p-values indicated above the corresponding bar.

The western blot analysis and the fluorescence levels of the eGFP-TMD chimeras suggested that none of them were toxic. Thus, the reduction in cell viability observed when they were transfected alongside BclxL and treated with doxorubicin was likely the result of their inhibition of BclxL and not a direct effect on cell viability. Nonetheless, we decided to test the viability of the cells after the transfection with eGFP-T20, eGFP-D1, eGFP-D2, or eGFP-xL. Our results indicated that neither D1 nor D2 was toxic to HeLa cells, a vital characteristic when designing a BclxL non-toxic inhibitor. (Figure 7e).

## Discussion

Virtually all pathological events associated with apoptosis deregulation can be linked to apoptosis resistance. Over-expression of anti-apoptotic Bcl2 members is common in newly diagnosed cancers and associated with resistance to several treatments, including various chemotherapeutic agents and γ-irradiation(54–57). Conversely, loss or down-regulation of pro-apoptotic members has been reported in many tumor types(58, 59).

Among the anti-apoptotic proteins, Bcl-2–like protein 1, better known as BclxL, displays relevant functions in several forms of cancer. In melanoma, BclxL participates in many of the hallmarks of this aggressive form of cancer, such as preventing cells from executing apoptosis and inducing drug resistance, cell migration and invasion, and angiogenesis(60). Because of the relevance of BclxL in the progression of cancer, different strategies have been considered to inhibit it(60). These strategies include antisense oligonucleotides, Proteolysis Targeting Chimeric (PROTAC) molecules, and BH3 mimetics. However, these approaches have not been successful, either because of their poor efficacy or the adverse effects associated with them.

Several methods have been described for the identification of TMD–TMD interactions, including *in vitro*, *in vivo*, and *in silico* approaches(61). To determine whether the TMD of BclxL interacts with other TMDs from the Bcl2 family, we chose BLaTM(19) and an adapted BiFC assay(24). The combination of these two methodologies allowed us to quantitatively analyze potential TMD–TMD interactions (BLaTM) and investigate their presence *in vivo* in eukaryotic cells (BiFC). Furthermore, the adapted BiFC assay replicated the native membrane topology of the proteins under study. Bcl2 proteins are characterized by the absence of a signal peptide and by a single TMD in their Ct end (40), this type of membrane proteins are known as tail-anchored proteins(43). By locating the TMD at the Ct end of our BiFC chimeras, we replicated the location/orientation and membrane targeting of a Bcl2 protein and, we inferred, the necessary features for TMD–TMD interaction.

Both the BLaTM and BiFC assays facilitated the analysis of each potential TMD interaction independently. There are many advantages of reductionist and isolating approaches. Nevertheless, there are some limitations to consider, including that the formation of higher-order oligomers (e.g., trimers, tetramers) could not be excluded. Additionally, although the formation of homo-oligomers is self-sufficient (no other TMD is involved), the formation of a hetero-oligomer might require prior assembly of an intramembrane homo-oligomer, and BiFC or BLaTM do not yield the relevant information on this matter.

The results obtained with the BiFC and BLaTM approaches were very similar, nonetheless, we observed some differences. These differences can largely be explained by discrepancies in the nature and location of reporters and, more important, in the membrane environment, prokaryotes vs. eukaryotes, which affects TMD–TMD interactions(24, 62–64). The disparity observed for the BclxL and Bax TMD-TMD interaction could also be attributed to the low expression levels of Bax TMD in 293T cells. Expression of the Bax TMD domain has been shown to induce some toxicity in eukaryotic cells(16, 18).

There have been described 3 modes in which BclxL could block apoptosis. Mode 0 suggests that BclxL shifts the equilibrium between soluble and membrane-bound Bax(49, 50). Mode 1 indicates that BclxL sequesters BH3 activators(65, 66). Lastly, Mode 2, proposes that BclxL directly sequester pro-apoptotic Bax(66–68). We detected interactions between the TMD of BclxL and the TMD of pro-apoptotic, anti-apoptotic, and BH3-only Bcl2 members. Furthermore, a membrane-bound BclxL chimera incapable of establishing intramembrane TMD-TMD interactions (such as Bclxl-T20) showed no anti-apoptotic properties. Of interest, the elimination of TMD–TMD interactions by substitution of the BclxL TMD with the TMD of mitochondrial Tomm20 protein had a stronger effect on BclxL inhibition of doxorubicin-induced apoptosis(27) than did the complete elimination of the TMD. Thus, while we have no data regarding Mode 0 for the induction of apoptosis, in membranes, both Mode 1 and 2 are compatible with our results. Furthermore, we provided data suggesting that in Mode 1 and/or 2 the interactions of the TMD of BclxL with other TMDs of the Bcl2 family members play a key role. Of note, *in vivo*, all the described TMD–TMD interactions could occur simultaneously or display a hierarchical order in which some interactions are preferred. Currently, data exist to support either possibility(69, 70), and the two are not mutually exclusive and most likely occur simultaneously in cells.

To block the intramembrane interactions of BclxL and consequently its anti-apoptotic function, we designed three potential inhibitors (D1, D2, and D3), based on a model of a putative BclxL TMD homodimer. Our data does not inform on the oligomeric state of BclxL TMD, however, there is data suggesting that BclxL forms homo-dimers both in solution and in the membrane(70). All three inhibitors’ designs retained the Gly residues within the TMD of BclxL (Gly8 and Gly13, D1) or substituted one of them with an Ala residue (Gly8Ala, D2 and D3). Small residues, particularly Gly, have been observed in other TMD–TMD oligomers (e.g., Gp A(20, 21), integrins(19), and even viral Bcl2 proteins(18)). The presence of these small residues in the designs facilitates helix-helix packing, maximizing the surface area and thus van der Waals forces, a major contributor to both homo- and hetero-intramembrane PPIs(71).

According to the results of the BLaTM assay, D2 forms a weaker interaction with the TMD of BclxL than does D1. The only difference between these two designs is in position 8, where there is a Gly in D1 and an Ala in D2. Our results (for both the BlaTM and the functional assays) indicate that a small structural difference, the substitution of Gly by Ala, has a major impact on the TMD–TMD interaction and the associated biological processes. These results, together with the substitutions at position 12, imply a high specificity in intramembrane interactions. Note that high specificity is necessary to achieve a complex interaction network like the one suggested by our TMD–TMD PPI scanning. However, it remains to be seen whether a single interaction can be interrupted specifically, which would indicate that each TMD–TMD PPI has its unique interaction profile. Most likely, given the reduced surface area of a TMD and the potentially large number of interaction partners, as is the case for the TMD of BclxL, some residues might be involved in interaction with multiple TMDs.

The expression of eGFP-D1 could block the anti-apoptotic function of BclxL. Likewise, D2 reduced the anti-apoptotic response of BclxL, although we found no statistically significant differences with the negative control. These results indicate that the strength of the interaction between D1 or D2 with the TMD of BclxL correlates with the inhibiting properties of the designs.

The subcellular localization by differential centrifugation in tandem with SWATH-MS suggests an organelle bound and a cytosolic form for BclxL (and Bax, see Supp. Information). It has been previously shown that Bcl2 family members shuttle between the mitochondria membrane and a soluble state(49–51). On the other hand, D1 behaves exclusively as an organelle bound protein. Thus, the D1-BclxL interaction and the corresponding inhibitory effect must occur in mitochondrial membranes. Any anti-cancer therapy based on our designs will require a vector that facilitates the insertion of the inhibitor into the MOM. Given the peptide nature of the inhibitors, there are two main possibilities for their delivery into cancer cells and their mitochondria. One is to use delivery systems for peptide-based drugs(72), such as cell-penetrating peptides (e.g., TAT), targeting peptides (e.g., RGD-4c), or stimuli-responsive peptides (e.g., pHLIP). The other is that instead of the peptide being delivered, a nucleic acid encoding the inhibitor (e.g., RNA/DNA lipid nanoparticles, polyplexes, or a viral vector) could be delivered. The hydrophobic nature of our designs complicates conventional peptide delivery, leading us to believe that the second option would be more appropriate. Of note, we could demonstrate that when eGFP-D1 is expressed from a DNA plasmid, it displays the same localization as BclxL in eukaryotic cells. This result indicates that the designs inherently can reach their active location when translated within cells with no further assistance.

In summary, we have provided evidence of the importance of TMD–TMD interactions in apoptosis control, particularly in the case of BclxL, and successfully designed, using a novel computational approach, sequences capable of specifically sequestering the TMD of BclxL. Through this sequestration capacity, our designs inhibit the anti-apoptotic action of BclxL. Our work shows a path to design effective inhibitors based only on the sequences of the target receptor. The fact that two of three designs exhibited the desired hetero and no homo-interactions highlights the accuracy of our strategy.

We hope that this work both advances our understanding of sequence-specific recognition in membranes and opens the way to a new generation of anti-cancer drugs without the limitations of current BclxL inhibitors.

## Materials and Methods

### Cell cultures, plasmids, and reagents

Human embryonic kidney 293T cells (HEK 293T), and human epithelial cervical cancer cells (HeLa) were cultured in Dulbecco’s modified Eagle’s medium (DMEM) (Gibco) supplemented with 10% fetal bovine serum (FBS) (Gibco), and penicillin-streptomycin (P/S) (100 U/mL) (Gibco). Cells were grown at 37 °C, with 5% CO_2_. Transfection of DNA into eukaryotic cells was performed in a serum-free medium (Gibco) with Lipofectamine 2000 (Invitrogen) according to the manufacturer’s specifications.

The TMD sequences were synthesized by Invitrogen (GeneArt gene synthesis), PCR amplified, and subcloned into the appropriated vector either using the InFusion cloning system following the manufacturer’s protocol (Takara). Mutations into the TMD were introduced by site-directed mutagenesis using the Quick Change II kit following the manufacturer’s instructions (Agilent Technologies). All DNA manipulations were confirmed by the sequencing of plasmid DNAs (Macrogen Spain). Transfection of DNA into eukaryotic cells was performed in Opti-MEM reduced serum medium (Gibco) with Lipofectamine 2000 (Invitrogen) according to the manufacturer’s specifications.

### Bimolecular fluorescent complementation (BiFC) assay

For the generation of BiFC chimeric plasmids including the Nt or Ct of the Venus Fluorescent Protein (VN, VC, respectively) plasmids were modified (Addgene #27097, #22011, a gift from Chang-Deng H) (73) to clone the cellular and viral Bcl2 TMDs at the Ct of the VFP(18). Chimeras (500 ng VN + 500 ng VC) were transfected into 2.10^5^ HEK 293T cells together with a plasmid expressing Renilla luciferase under the CMV promoter (pRL-CMV) (50 ng) for signal normalization. Cells were incubated at 37 °C, 5% CO_2_ for 48h, PBS washed, and collected for fluorescence and luciferase measurements (Victor X3 plate reader). For the Renilla luciferase readings, we used the Renilla Luciferase Glow Assay Kit (Pierce, Thermofisher) according to the manufacturer’s protocol. In each experiment, the fluorescence/luminescence ratio obtained with the GpA homodimer was used as a 100% oligomerization value and the rest of the values were adjusted accordingly. All experiments were done at least in triplicates.

### BLaTM assay

Competent E. coli BL21-DE3 cells were co-transformed with N-BLa and C-BLa plasmids, version 1.1(19), containing a given TMD pair and grown overnight at 37 °C on LB-agar plates containing 34 μg/mL of chloramphenicol (Cm) and 35 μg/mL of kanamycin (Kan) for plasmid inheritance. After o/n incubation at 37 °C, colonies were either picked for immediate use or the plates were sealed with Parafilm (Pechiney Plastic Packaging) and stored at 4 °C for up to one week. Overnight cultures were conducted by inoculating 5 mL of LB-medium (Cm, Kan) with 10 colonies from one agar plate, followed by o/n incubation in an orbital incubator at 37 °C, 200 rpm. An expression culture was started with a 1:10 dilution of the overnight culture in 4 mL expression medium: LB-medium (Cm, Kan) containing 1.33 mM arabinose. After 4 h at 37 °C, the expression cultures were diluted to an OD_600_ = 0.1 in the expression medium. To expose the bacteria to different ampicillin concentrations, an LD_50_ culture was prepared by pipetting 100 μL of the diluted expression culture into each cavity of a 96-deep well plate (96 square well, 2 mL, VWR) containing 400 μL of expression media (final OD_600_ = 0.02). Freshly prepared ampicillin stock (100 mg/mL in ethanol) was added, resulting in ampicillin concentrations ranging from 0 to 350 μg/mL, depending on the affinity of the TMD under investigation. As a rule, the maximum ampicillin concentration to be used for a particular case should be about twice the mean LD_50_. The plates were incubated in a moisturized container for 16 h at 37 °C and 250 rpm on a shaker (shaking amplitude 10 mm, KS 260 Basic, IKA) containing tips in every well to ensure proper agitation. Cell density was measured via absorbance at 544 nm in a microplate reader (Victor X3, Perkin Elmer). To minimize clonal variation, at least two transformations were done and at least two separate LD_50_ cultures were inoculated from each batch of transformed bacteria using ten colonies for each culture. Thus, at least 40 colonies entered each determination of LD_50_. To measure and collect LD_50_ values from the dose-response curves, we used Prism 9 from GraphPad.

To analyze the expression levels of the chimeras, competent E. coli BL21-DE3 cells were transformed with one N-BLa or C-BLa plasmid, version 1.1, containing a given TMD and grown overnight at 37 °C on LB-agar plates containing 34 μg/mL of Cm (for N-BLa) or 35 μg/mL of Kan (for C-BLa) for plasmid inheritance. After o/n incubation at 37 °C, cultures were conducted by inoculating 5 mL of LB-medium (Cm or Kan) with 10 colonies from one agar plate, followed by o/n incubation in an orbital incubator at 37 °C, 200 rpm. An expression culture was started with a 1:10 dilution from the overnight culture in 4 mL expression medium: LB-medium (Cm or Kan) containing 1.33 mM arabinose. After 4 h at 37 °C, the expression cultures were diluted to an OD600 = 0.1 in 5 ml of expression medium (final volume) and grown o/n, 37 °C, 200xg. The morning after, 100 μL from each culture were transferred to a black 96-well plate to measure the fluorescence (Victor X3, Perkin Elmer).

### In vitro transcription and translation

The Lep-derived constructs were assayed using the TNT T7 Quick Coupled System (#L1170, Promega). Each reaction containing 1 µL of PCR product, 0.5 µL of EasyTag™ EXPRESS 35S Protein Labeling Mix (Perkin Elmer) (5.5 µCi), and 0.3 µL of microsomes (tRNA Probes) was incubated at 30 °C for 90 min. Samples were analyzed by SDS-PAGE. The bands were quantified using a Fuji FLA-3000 phosphoimager and the Image Reader 8.1 software. Free energy was calculated using: ΔGapp = −RT.lnKapp, where Kapp = f2g/f1g being f1g and f2g the fraction of single glycosylated and double glycosylated protein, respectively. Endoglycosidase H treatment (Roche) was carried out according to the specifications of the manufacturer.

### Cell-viability assays

To measure doxorubicin-induced apoptosis 1.5 × 10^6^ HeLa cells were plated in a 24 wells plate containing 0.5 ml of media in each well. After overnight incubation, each well was transfected in triplicates with 500 ng of DNA. After 24 h of expression, cells were treated with doxorubicin (stock 2 mM in DMSO) achieving a final concentration of 15 µM. Approximately, 16 h post-treatment cells (including those in the supernatant) were collected and their viability was measured using Trypan blue and using an automated cell counter (Invitrogen, Countess™ II). At least, 2 measurements per well were done. Alternatively, 16/24 h post-treatment media was removed and the cells were washed (PBS x2) and fixed with 6% Glutaraldehyde containing 0.5% of crystal violet. After 20 min fixed cells were washed with water (x3) before visualization.

To measure the anti-BclxL effect of the designs, 1.5 × 10^6^ HeLa cells were plated in a 24-well plate (0.5 ml of media in each well). After overnight incubation, cells were transfected (in triplicate) with 500 ng of either eGFP-XL, eGFP-T20, eGFP-D1, eGFP-D2, or empty pCAGGS (negative control). Additionally, cells were co-transfected with 500 ng of BclxL full length except for the negative control where 500 ng of empty pCAGGS were transfected. After 24 h of expression, cells were treated with doxorubicin (stock 2 mM in DMSO) achieving a final concentration of 15 µM. Approximately, 16 h post-treatment cells (including those in the supernatant) were collected and their viability was measured using Trypan blue and using an automated cell counter (Invitrogen, Countess™ II). At least, 2 measurements per well were done.

To study the toxicity of individual chimeras, 1.5 × 10^6^ HeLa cells were plated in a 24 wells plate containing 0.5 ml of media in each well. After overnight incubation, each well was transfected in triplicates with 500 ng of a plasmid containing eGFP-T20, eGFP-D1, eGFP-D2, or eGFP-xL. 24 h post-transfection, cells were collected (including those in the supernatant) and their viability was measured using Trypan blue and using an automated cell counter (Invitrogen, Countess™ II). At least, 2 measurements per well were done.

To analyze the expression levels of the chimeras, 1.5 x 10^6^ HeLa cells were plated in a 24 wells plate containing 0.5 ml of media in each well. After overnight incubation, each well was transfected in duplicates with 500 ng of a plasmid containing eGFP-T20, eGFP-D1, eGFP-D2, or eGFP-xL. Cells were incubated at 37 °C, 5% CO2 for 24 h, PBS washed, and collected for fluorescence measurements (Victor X3 plate reader) or western blot analysis.

### Inhibitor design

The design of the inhibitors started with the modeling of the BclxL – BclxL homo-interaction. The modeling was done using TMHOP (Trans-membrane Homo Oligomer Predictor; https://TMHOP.weizmann.ac.il)(5). The lowest energy models were selected and visually inspected. Using the selected model as the starting template, the design step was prepared using FuncLib(31) with the membrane filter by selecting mutations that obeyed the following rules: 1. Positive design for a heterodimer (Non-symmetric FuncLib; ΔΔG<+1 Rosetta energy units; R.e.u.); 2. Negative selection for a homodimer (symmetric FuncLib; ΔΔG>+5 R.e.u.). The resulting TMDs were manually selected after visualizing the predicted structures.

### TMD interaction surface area and TMD-TMD crossing angle calculation

Amino acid residues in the interface were selected using InterfaceResidues.py script by Jason Vertrees (pymolwiki.org/index.php/InterfaceResidues). Then, the surface area in square Angstroms was calculated using get_area command in The PyMOL (TM) Molecular Graphics System, Version 2.3.0 Schrodinger, LLC for the complete TMD and the selected interface area. The crossing angle between helices was calculated using AngleBetweenHelices.py script by Thomas Holder (pymolwiki.org/index.php/AngleBetweenHelices).

### Subcellular protein localization by differential ultracentrifugation

The proteins of interest were expressed in the Hek-293T human cell line (37 °C and 5% CO_2_ with DMEM supplemented with FBS, Penstrep, and Amphotericin B). For each construct, two plates with 7 x 10^6^ cells each were seeded. After 24 hours cells were transfected with plasmids encoding the BclXL or eGFP-D1 using PEI. After 24 hours cells were harvested, washed with PBS buffer, and collected by trypsinization. Next, cells were centrifugated (200g 5min) and resuspended with 1.5 mL of lysis buffer (0.25 M sucrose, 10 mM HEPES pH 7.4, 2 mM EDTA, 2 mM magnesium acetate, cOmplete protease inhibitor (Roche)). Next cells were lysed in an Isobiotech homogenizer using the 12 μm clearance size ball. Cells were passed through the homogenizer until 90% of cell disruption was achieved followed by centrifugation 2×200g 5min to eliminate intact cells. The supernatant was collected and a series of differential centrifugations were carried out at 4°C. Precisely, 1.000 g 10 min, 3.000g 10 min, 5.000 g 10 min, 9,000 g 15 min, 12.000 g 15 min, 15.000 g 15 min, 30.000 g 20 min, 79.000 g 43 min, 120.000 g 45 min. Pellets were resuspended (8 M urea, 0.15% SDS, 50 mM HEPES pH 8.5) and the protein concentration was determined by Bradford. All samples were submitted for SWATH-MS quantification at the Proteomics core facility of the University of Valencia. Data analysis was performed using the R Bioconductor(47, 74) packages MSnbase v2.6.1(75) and pRoloc v1.21.9(76) as described previously(47).

### Confocal microscopy

Confocal micrographs were done at the Microscopy Core Facility of the SCSIE (University of Valencia) using an Olympus FV1000 confocal microscope with a ×60 oil lens. Mitochondria mCherry fluorescent-labeled marker (mCherry-Mito) was obtained from Addgene plasmid repository #55146, a gift from Michael Davidson, Institute of Molecular Biophysics and Center for Materials Research and Technology, The Florida State University. HeLa cells (5 × 10^3^ cells/well) were seeded on 10 mm cover slides treated with poly-Lys and placed in 24-well plates. The next day, cells were transfected with the appropriate plasmids. After 24 h, the cells were fixed (4% paraformaldehyde) and DAPI stained before image capture. A 1:1000 dilution in TBS 0.005 % Tween Rabbit anti-c-Myc (Sigma PLA0001) antibody followed by an anti-Rabbit Alexa 488 conjugated (Life Technologies A21206) (1:1000) was used to label BclxL FL, BclxL T20, and BclxL ΔTM proteins. For the eGFP chimeras: eGFP-xL, eGFP-D1, eGFP-D1, and eGFP-D2 no antibody was needed as it was possible to use the eGFP itself. Pictures were taken in an Olympus FV1000 confocal microscope. Laser intensity was individually adjusted in all samples. Pictures were not used for quantification.

### Western blot analysis

Cell monolayers were lysed in Laemmlís sample buffer (62.5 mM Tris-HCl [pH 6.8], 2 % sodium dodecyl sulfate [SDS], 0.01% bromophenol blue, 10 % glycerol and 5 % β-mercaptoethanol). Protein samples were subjected to 12 % SDS-polyacrylamide gel electrophoresis (PAGE) and transferred to nitrocellulose membranes (BioRad). Membranes were blocked for 30 min at room temperature in Tris-buffered saline supplemented with 0.05 % Tween 20 (TBS-T) containing 5 % non-fat dry milk and later incubated with primary antibodies diluted in the same buffer at 4 °C overnight. Antibodies used in this study were β-actin (Santa Cruz Biotechnology SC-47778), c-Myc (Sigma PLA0001 or Roche 11667149001), Histone 3 (Sigma H0164), and Flag (Sigma B3111). Then, membranes were washed with TBS-T and incubated with goat anti-mouse IgG-peroxidase conjugate (Sigma DC02L) for 1 h at room temperature and washed again. All antibodies were used at a 1:10,000 dilution in TBS-T with 5 % non-fat dry milk. Detection of immunoreactive proteins was carried out using the enhanced chemiluminescence (ECL) reaction (SuperSignal ThermoScientific) and detected by the ChemiDoc Touch Imaging System (BioRad).

## Acknowledgments

We thank the Generalitat Valenciana (PROMETEO/2019/065) and the Spanish Ministry of Science and Innovation (PID2020-119111GB-I00). G.-D. is the recipient of a predoctoral grant from the Spanish Ministry of Universities (FPU18/05771). The Fleishman lab was supported by the Dr. Barry Sherman Institute for Medicinal Chemistry and by a charitable donation in memory of Sam Switzer. We thank P. Selvi for excellent technical assistance.

## References

1. L. Cao, et al., Design of protein binding proteins from target structure alone. Nature (2022) https://doi.org/10.1038/s41586-022-04654-9.

2. Y. Bouchiba, M. Ruffini, T. Schiex, S. Barbe, Computational Design of Miniprotein Binders. Methods Mol. Biol. Clifton NJ 2405, 361–382 (2022).

3. M. Delaunay, T. Ha-Duong, Computational Tools and Strategies to Develop Peptide-Based Inhibitors of Protein-Protein Interactions. Methods Mol. Biol. Clifton NJ 2405, 205–230 (2022).

4. H. Yin, et al., Computational design of peptides that target transmembrane helices. Science 315, 1817–1822 (2007).

5. J. Y. Weinstein, A. Elazar, S. J. Fleishman, A lipophilicity-based energy function for membrane-protein modelling and design. PLoS Comput. Biol. 15 (2019).

6. A. Elazar, et al., Mutational scanning reveals the determinants of protein insertion and association energetics in the plasma membrane. eLife 5, e12125 (2016).

7. A. Elazar, J. J. Weinstein, J. Prilusky, S. J. Fleishman, Interplay between hydrophobicity and the positive-inside rule in determining membrane-protein topology. Proc. Natl. Acad. Sci. U. S. A. 113, 10340–10345 (2016).

8. A. Elazar, et al., De novo-designed transmembrane domains tune engineered receptor functions. eLife 11, e75660 (2022).

9. M. O. Hengartner, The biochemistry of apoptosis. Nature 407, 770–776 (2000).

10. H. Kim, et al., Hierarchical regulation of mitochondrion-dependent apoptosis by BCL-2 subfamilies. Nat. Cell Biol. 8, 1348–1358 (2006).

11. Z. N. Oltvai, C. L. Milliman, S. J. Korsmeyer, Bcl-2 heterodimerizes in vivo with a conserved homolog, Bax, that accelerates programmed cell death. Cell 74, 609–619 (1993).

12. K. Wang, X. M. Yin, D. T. Chao, C. L. Milliman, S. J. Korsmeyer, BID: a novel BH3 domain-only death agonist. Genes Dev. 10, 2859–2869 (1996).

13. A. Kelekar, B. S. Chang, J. E. Harlan, S. W. Fesik, C. B. Thompson, Bad is a BH3 domain-containing protein that forms an inactivating dimer with Bcl-XL. Mol. Cell. Biol. 17, 7040–7046 (1997).

14. K. Cosentino, A. J. García-Sáez, Bax and Bak Pores: Are We Closing the Circle? Trends Cell Biol. 27, 266–275 (2017).

15. S. Dadsena, L. E. King, A. J. García-Sáez, Apoptosis regulation at the mitochondria membrane level. Biochim. Biophys. Acta BBA - Biomembr. 1863, 183716 (2021).

16. V. Andreu-Fernández, et al., Bax transmembrane domain interacts with prosurvival Bcl-2 proteins in biological membranes. Proc. Natl. Acad. Sci. U. S. A. 114, 310–315 (2017).

17. E. Lucendo, et al., Mcl-1 and Bok transmembrane domains: Unexpected players in the modulation of apoptosis. Proc. Natl. Acad. Sci. U. S. A. 117, 27980–27988 (2020).

18. M. J. García-Murria, et al., Viral Bcl2s’ transmembrane domain interact with host Bcl2 proteins to control cellular apoptosis. Nat. Commun. 11, 6056 (2020).

19. C. Schanzenbach, F. C. Schmidt, P. Breckner, M. G. Teese, D. Langosch, Identifying ionic interactions within a membrane using BLaTM, a genetic tool to measure homo- and heterotypic transmembrane helix-helix interactions. Sci. Rep. 7, 43476 (2017).

20. M. A. Lemmon, J. M. Flanagan, H. R. Treutlein, J. Zhang, D. M. Engelman, Sequence specificity in the dimerization of transmembrane alpha-helices. Biochemistry 31, 12719–12725 (1992).

21. K. R. MacKenzie, J. H. Prestegard, D. M. Engelman, A transmembrane helix dimer: structure and implications. Science 276, 131–133 (1997).

22. M. Orzáez, E. Pérez-Payá, I. Mingarro, Influence of the C-terminus of the glycophorin A transmembrane fragment on the dimerization process. Protein Sci. Publ. Protein Soc. 9, 1246– 1253 (2000).

23. T. K. Kerppola, Design and implementation of bimolecular fluorescence complementation (BiFC) assays for the visualization of protein interactions in living cells. Nat. Protoc. 1, 1278–1286 (2006).

24. B. Grau, et al., The role of hydrophobic matching on transmembrane helix packing in cells. Cell Stress 1, 90–106 (2017).

25. R. W. Rooswinkel, et al., Antiapoptotic potency of Bcl-2 proteins primarily relies on their stability, not binding selectivity. Blood 123, 2806–2815 (2014).

26. M. González-García, et al., bcl-XL is the major bcl-x mRNA form expressed during murine development and its product localizes to mitochondria. Dev. Camb. Engl. 120, 3033–3042 (1994).

27. W. Fang, J. J. Rivard, D. L. Mueller, T. W. Behrens, Cloning and molecular characterization of mouse bcl-x in B and T lymphocytes. J. Immunol. 153, 4388–4398 (1994).

28. N. Zamzami, C. Brenner, I. Marzo, S. A. Susin, G. Kroemer, Subcellular and submitochondrial mode of action of Bcl-2-like oncoproteins. Oncogene 16, 2265–2282 (1998).

29. L. Martínez-Gil, A. Saurí, M. A. Marti-Renom, I. Mingarro, Membrane protein integration into the endoplasmic reticulum. FEBS J. 278, 3846–3858 (2011).

30. U. S. Chio, H. Cho, S.-O. Shan, Mechanisms of Tail-Anchored Membrane Protein Targeting and Insertion. Annu. Rev. Cell Dev. Biol. 33, 417–438 (2017).

31. O. Khersonsky, et al., Automated Design of Efficient and Functionally Diverse Enzyme Repertoires. Mol. Cell 72, 178–186.e5 (2018).

32. R. Netzer, et al., Ultrahigh specificity in a network of computationally designed protein-interaction pairs. Nat. Commun. 9, 5286 (2018).

33. C. Bolenz, et al., Optimizing chemotherapy for transitional cell carcinoma by application of bcl-2 and bcl-xL antisense oligodeoxynucleotides. Urol. Oncol. 25, 476–482 (2007).

34. L. Martínez-Gil, et al., Membrane integration of poliovirus 2B viroporin. J. Virol. 85, 11315– 11324 (2011).

35. M. Bañó-Polo, C. A. Martínez-Garay, B. Grau, L. Martínez-Gil, I. Mingarro, Membrane insertion and topology of the translocon-associated protein (TRAP) gamma subunit. Biochim. Biophys. Acta BBA - Biomembr. 1859, 903–909 (2017).

36. C. Lundin, H. Kim, I. Nilsson, S. H. White, G. von Heijne, Molecular code for protein insertion in the endoplasmic reticulum membrane is similar for N(in)-C(out) and N(out)-C(in) transmembrane helices. Proc. Natl. Acad. Sci. U. S. A. 105, 15702–15707 (2008).

37. L. Kasturi, H. Chen, S. H. Shakin-Eshleman, Regulation of N-linked core glycosylation: use of a site-directed mutagenesis approach to identify Asn-Xaa-Ser/Thr sequons that are poor oligosaccharide acceptors. Biochem. J. 323 (Pt 2), 415–419 (1997).

38. I. Mingarro, G. von Heijne, P. Whitley, Membrane-protein engineering. Trends Biotechnol. 15, 432–437 (1997).

39. K. Braunger, et al., Structural basis for coupling protein transport and N-glycosylation at the mammalian endoplasmic reticulum. Science 360, 215–219 (2018).

40. V. Andreu-Fernández, et al., The C-terminal Domains of Apoptotic BH3-only Proteins Mediate Their Insertion into Distinct Biological Membranes. J. Biol. Chem. 291, 25207–25216 (2016).

41. T. Hessa, et al., Molecular code for transmembrane-helix recognition by the Sec61 translocon. Nature 450, 1026–1030 (2007).

42. T. Hessa, et al., Recognition of transmembrane helices by the endoplasmic reticulum translocon. Nature 433, 377–381 (2005).

43. R. S. Hegde, R. J. Keenan, The mechanisms of integral membrane protein biogenesis. Nat. Rev. Mol. Cell Biol. 23, 107–124 (2022).

44. A. Geladaki, et al., Combining LOPIT with differential ultracentrifugation for high-resolution spatial proteomics. Nat. Commun. 10, 331 (2019).

45. F. Zhang, W. Ge, G. Ruan, X. Cai, T. Guo, Data-Independent Acquisition Mass Spectrometry-Based Proteomics and Software Tools: A Glimpse in 2020. Proteomics 20, e1900276 (2020).

46. R. J. Rotello, T. D. Veenstra, Mass Spectrometry Techniques: Principles and Practices for Quantitative Proteomics. Curr. Protein Pept. Sci. 22, 121–133 (2021).

47. L. M. Breckels, C. M. Mulvey, K. S. Lilley, L. Gatto, A Bioconductor workflow for processing and analysing spatial proteomics data. F1000Research 5, 2926 (2016).

48. O. M. Crook, C. M. Mulvey, P. D. W. Kirk, K. S. Lilley, L. Gatto, A Bayesian mixture modelling approach for spatial proteomics. PLoS Comput. Biol. 14, e1006516 (2018).

49. B. Schellenberg, et al., Bax exists in a dynamic equilibrium between the cytosol and mitochondria to control apoptotic priming. Mol. Cell 49, 959–971 (2013).

50. F. Edlich, et al., Bcl-x(L) retrotranslocates Bax from the mitochondria into the cytosol. Cell 145, 104–116 (2011).

51. R. Eskes, S. Desagher, B. Antonsson, J. C. Martinou, Bid induces the oligomerization and insertion of Bax into the outer mitochondrial membrane. Mol. Cell. Biol. 20, 929–935 (2000).

52. P. J. Thul, et al., A subcellular map of the human proteome. Science 356, eaal3321 (2017).

53. A. Christoforou, et al., A draft map of the mouse pluripotent stem cell spatial proteome. Nat. Commun. 7, 9992 (2016).

54. S. H. Kaufmann, et al., Elevated Expression of the Apoptotic Regulator Mcl-1 at the Time of Leukemic Relapse. Blood 91, 991–1000 (1998).

55. J. C. Reed, Regulation of apoptosis by bcl-2 family proteins and its role in cancer and chemoresistance. Curr. Opin. Oncol. 7, 541–546 (1995).

56. J. Lotem, L. Sachs, Control of apoptosis in hematopoiesis and leukemia by cytokines, tumor suppressor and oncogenes. Leukemia 10, 925–931 (1996).

57. P. W. Mesner, I. I. Budihardjo, S. H. Kaufmann, Chemotherapy-induced apoptosis. Adv. Pharmacol. San Diego Calif 41, 461–499 (1997).

58. S. Hafezi, M. Rahmani, Targeting BCL-2 in Cancer: Advances, Challenges, and Perspectives. Cancers 13, 1292 (2021).

59. P. D. Bhola, A. Letai, Mitochondria-Judges and Executioners of Cell Death Sentences. Mol. Cell 61, 695–704 (2016).

60. A. M. Lucianò, A. B. Pérez-Oliva, V. Mulero, D. Del Bufalo, Bcl-xL: A Focus on Melanoma Pathobiology. Int. J. Mol. Sci. 22, 2777 (2021).

61. G. Duart, B. Grau, I. Mingarro, L. Martinez-Gil, Methodological approaches for the analysis of transmembrane domain interactions: A systematic review. Biochim. Biophys. Acta Biomembr. 1863, 183712 (2021).

62. H. I. Petrache, A. Grossfield, K. R. MacKenzie, D. M. Engelman, T. B. Woolf, Modulation of glycophorin A transmembrane helix interactions by lipid bilayers: molecular dynamics calculations. J. Mol. Biol. 302, 727–746 (2000).

63. N. Flinner, O. Mirus, E. Schleiff, The influence of fatty acids on the GpA dimer interface by coarse-grained molecular dynamics simulation. Int. J. Mol. Sci. 15, 14247–14268 (2014).

64. L. Janosi, A. Prakash, M. Doxastakis, Lipid-modulated sequence-specific association of glycophorin A in membranes. Biophys. J. 99, 284–292 (2010).

65. A. J. García-Sáez, J. Ries, M. Orzáez, E. Pérez-Payà, P. Schwille, Membrane promotes tBID interaction with BCL(XL). Nat. Struct. Mol. Biol. 16, 1178–1185 (2009).

66. F. Llambi, et al., A unified model of mammalian BCL-2 protein family interactions at the mitochondria. Mol. Cell 44, 517–531 (2011).

67. Y. T. Hsu, R. J. Youle, Nonionic detergents induce dimerization among members of the Bcl-2 family. J. Biol. Chem. 272, 13829–13834 (1997).

68. E. H. Cheng, B. Levine, L. H. Boise, C. B. Thompson, J. M. Hardwick, Bax-independent inhibition of apoptosis by Bcl-XL. Nature 379, 554–556 (1996).

69. L. Fitzsimmons, et al., EBV BCL-2 homologue BHRF1 drives chemoresistance and lymphomagenesis by inhibiting multiple cellular pro-apoptotic proteins. Cell Death Differ. 27, 1554–1568 (2020).

70. S. Bleicken, A. Hantusch, K. K. Das, T. Frickey, A. J. Garcia-Saez, Quantitative interactome of a membrane Bcl-2 network identifies a hierarchy of complexes for apoptosis regulation. Nat. Commun. 8, 73 (2017).

71. L. Martinez-Gil, I. Mingarro, Viroporins, Examples of the Two-Stage Membrane Protein Folding Model. Viruses 7, 3462–3482 (2015).

72. D. Berillo, A. Yeskendir, Z. Zharkinbekov, K. Raziyeva, A. Saparov, Peptide-Based Drug Delivery Systems. Medicina (Mex.) 57, 1209 (2021).

73. Y. Kodama, C.-D. Hu, An improved bimolecular fluorescence complementation assay with a high signal-to-noise ratio. BioTechniques 49, 793–805 (2010).

74. R. C. Gentleman, et al., Bioconductor: open software development for computational biology and bioinformatics. Genome Biol. 5, R80 (2004).

75. L. Gatto, K. S. Lilley, MSnbase-an R/Bioconductor package for isobaric tagged mass spectrometry data visualization, processing and quantitation. Bioinforma. Oxf. Engl. 28, 288–289 (2012).

76. L. Gatto, L. M. Breckels, S. Wieczorek, T. Burger, K. S. Lilley, Mass-spectrometry-based spatial proteomics data analysis using pRoloc and pRolocdata. Bioinforma. Oxf. Engl. 30, 1322–1324 (2014).

