## Supplemental Figures for "Computational design of BclxL inhibitors that target transmembrane domain interactions"

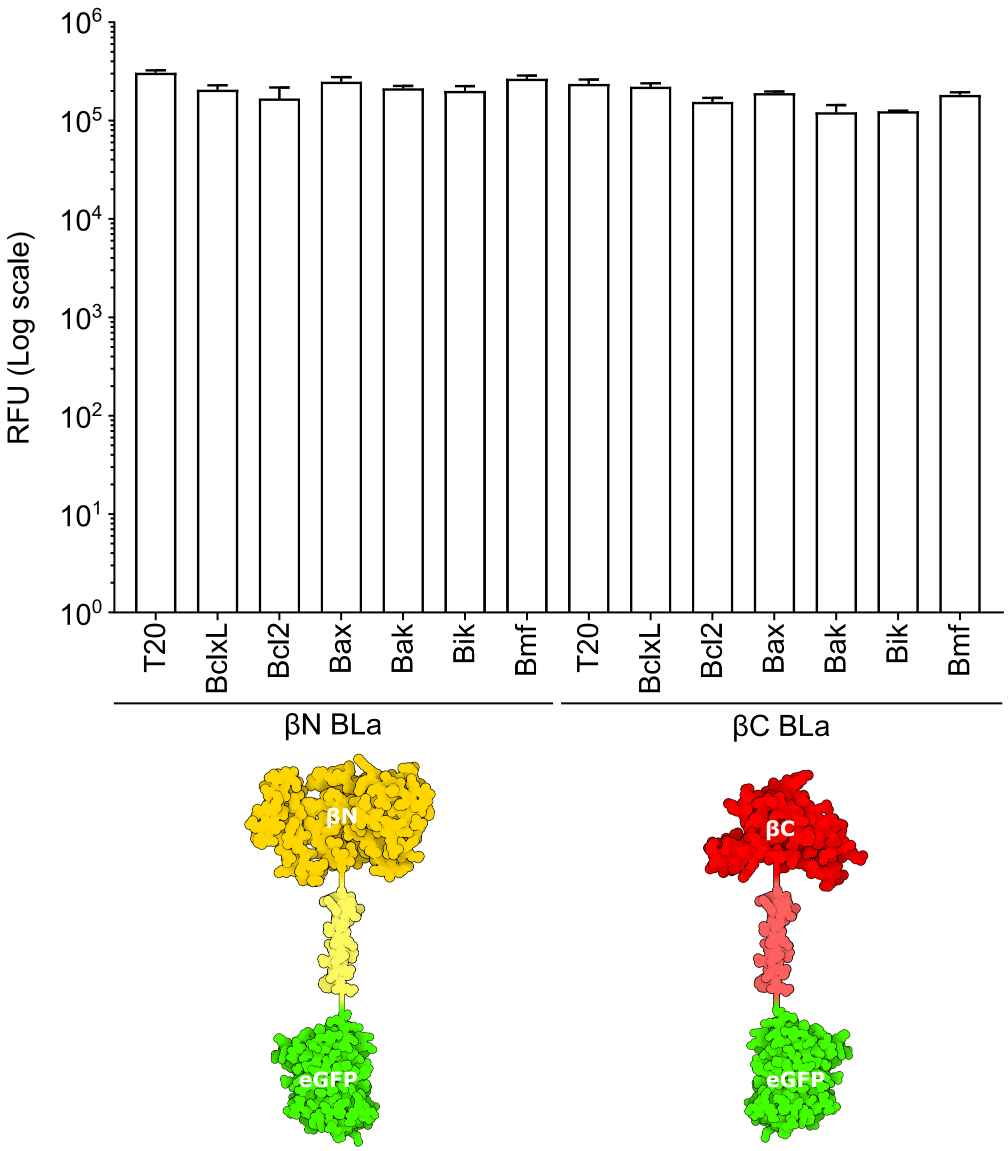


**Figure S1. Analysis of the expression levels of the BLaTM chimeras.** The eGFP-associated relative fluorescence ($\lambda_{exc}$ 485 nm, $\lambda_{em}$ 535nm) values for all BLaTM chimeras are shown. The mean and standard deviation of three independent experiments are shown (n = 3).

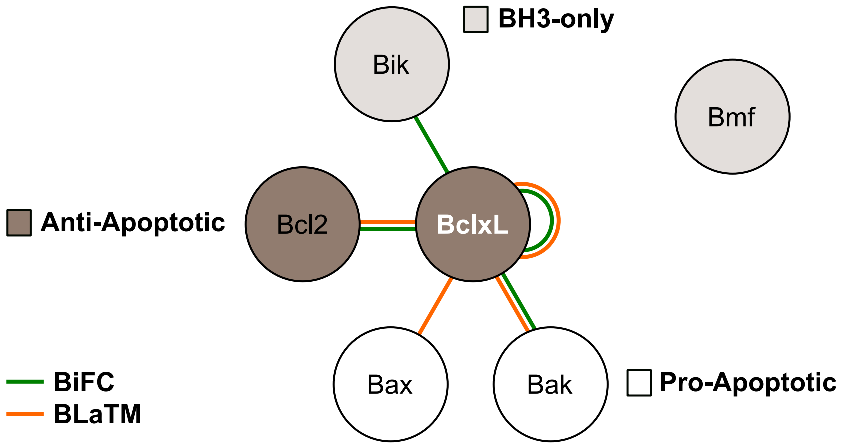


**Figure S2. TMD-TMD interaction network of BclxL.** A network representation of BclxL TMD interactions. The figure includes the results of the BiFC (green lines) and BLaTM assays (orange lines). Solid lines represent interactions, while TMDs are represented by nodes. Anti-apoptotic, pro-apoptotic, and BH3-only nodes are colored distinctively.

a

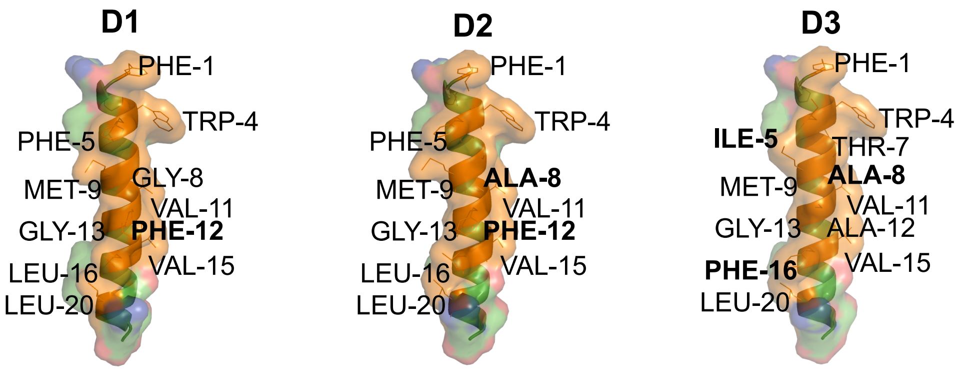


b

| Dimer | Name | Sequence | Tot. surface (Å²) | Inter. surface (Å²) | Angle (º) |
|---|---|---|---|---|---|
| xL-xL | BclxL | FNRWFLTGMTVAGVVLLGSLFSR | 2389.487 | 1244.083 | 48.00 |
| | BclxL | FNRWFLTGMTVAGVVLLGSLFSR | 2389.487 | 1244.083 | |
| xL-D1 | BclxL | FNRWFLTGMTVAGVVLLGSLFSR | 2389.487 | 1220.531 | 50.07 |
| | D1 | FNRWFLTGMTVFGVVLLGSLFSR | 2434.134 | 1206.933 | |
| xL-D2 | BclxL | FNRWFLTGMTVAGVVLLGSLFSR | 2389.487 | 1220.531 | 50.08 |
| | D2 | FNRWFLTAMTVFGVVLLGSLFSR | 2452.863 | 1223.097 | |
| xL-D3 | BclxL | FNRWFLTGMTVAGVVLLGSLFSR | 2389.487 | 1134.406 | 49.41 |
| | D3 | FNRWILTAMTVAGVVFLGSLFSR | 2453.863 | 1244.083 | |

**Figure S3. Interaction between the TMD of BclxL and D1, D2, and D3 designs. a**. Structural representations of D1, D2, and D3 designs. Numbers correspond to the position of the amino acids in the design. Differences among the designs are highlighted in bold. **b**. The amino acids on the interaction surfaces of the potential homo and heterodimers are marked in orange. Changes in the designs over the TMD sequence of BclxL are highlighted in yellow. The total surface area of each monomer and the area buried in the interaction with the corresponding TMD in a potential dimer are shown (Å$^2$). Additionally, the crossing angle of the monomers for each interaction is indicated.

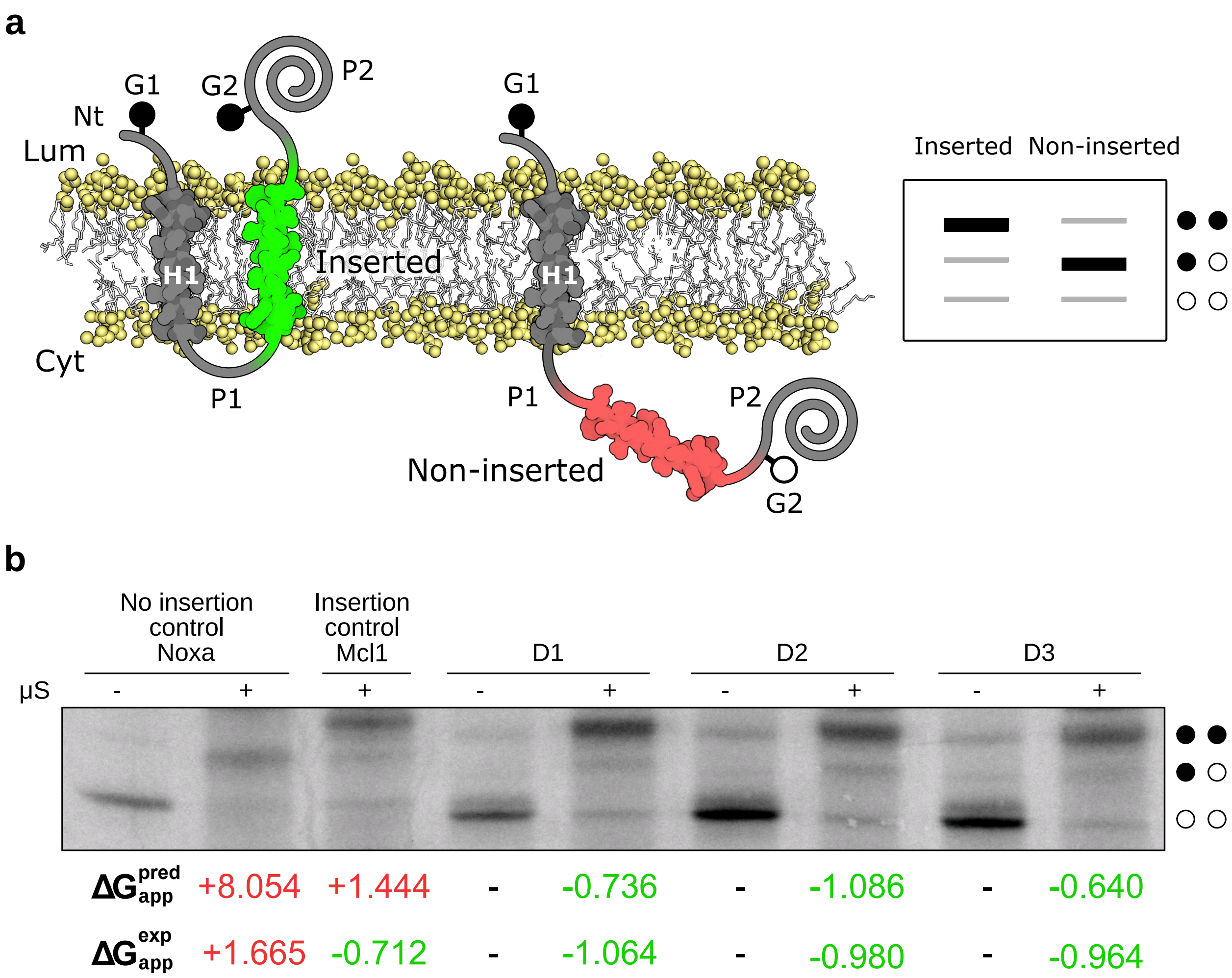


**Figure S4. Insertion of D1, D2, and D3 in eukaryotic membranes. a**. Schematic representation of the leader peptidase (Lep) model protein. G1 and G2 denote artificial glycosylation acceptor sites. The sequence under investigation is introduced between the P1 and P2 domains (replacing the natural occuring H2 domain, not shown) of Lep. Recognition of the tested sequence by the translocon machinery as a TMD (highlighted in green) results in the modification of the G1 and G2 acceptor sites. In contrast, only G1 will be glycosylated if the sequence being tested is not recognized as a TMD (shown in red) and thus not inserted into ER-derived membranes. The glycosylation state of a protein can be monitored by SDS-PAGE based on the increase in molecular weight associated with the addition of a sugar moiety. A mock SDS-PAGE is represented on the right. The absence of glycosylation of G1 and G2 acceptor sites is indicated by two white dots, single glycosylation by one white and one black dot, and double glycosylation by two black dots. **b**. A representative example (n = 3) of in vitro protein translation in the presence (+) or absence (−) of ER-derived microsomes (μS). The absence of glycosylation of G1 and G2 acceptor sites is indicated by two white dots, single glycosylation by one white and one black dot, and double glycosylation by two black dots. Additionally, the predicted and experimental ΔGs for the insertion in kcal/mol ($\Delta G_{app}^{pred}$ and $\Delta G_{app}^{exp}$) are presented below the image. Experimental results are the average of at least three independent experiments. Green numbers indicate negative ΔGs (insertion of the tested sequence), and red numbers indicate positive ΔGs (no insertion of the tested sequence).

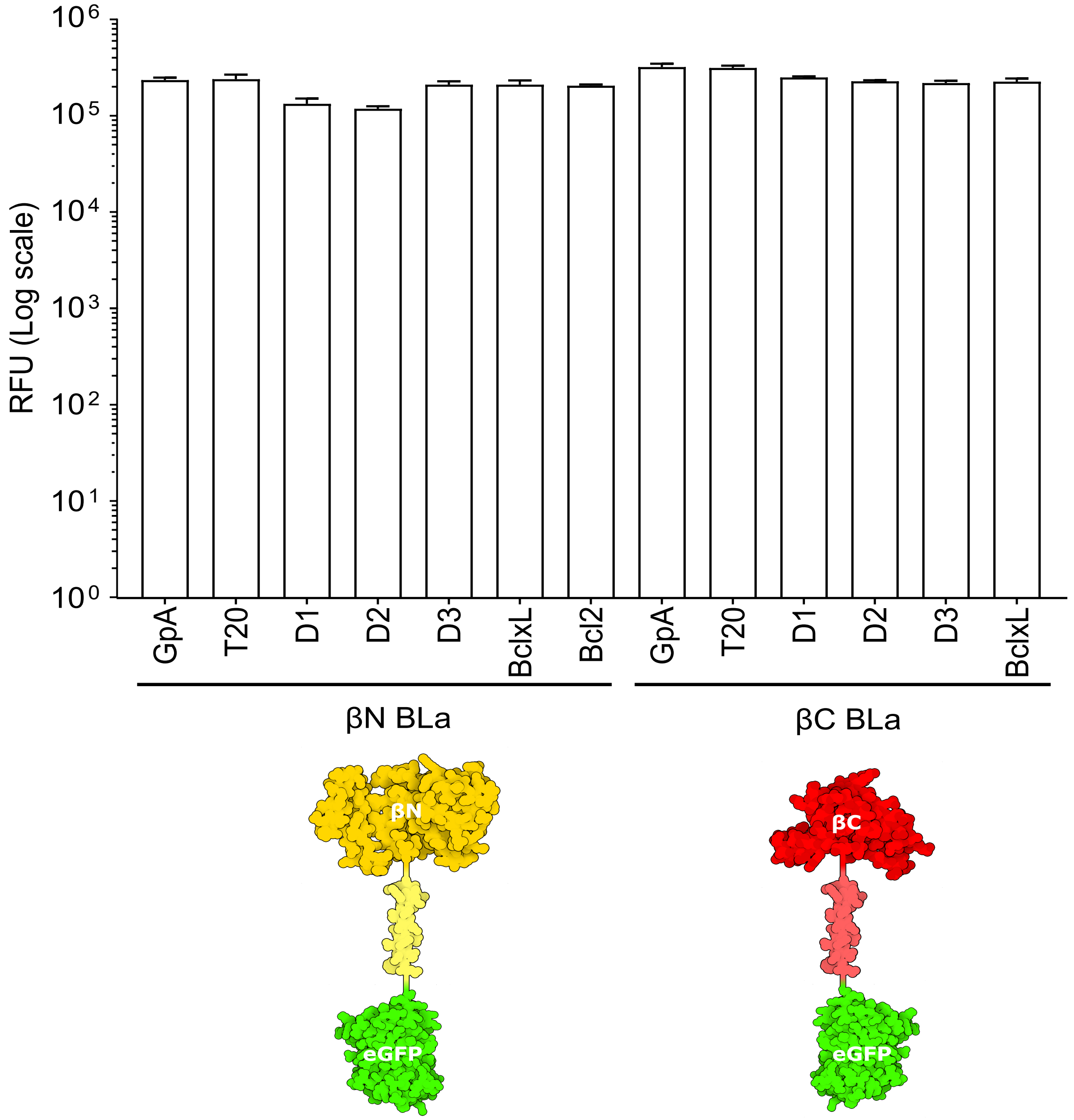


**Figure S5. Analysis of TMD—TMD interactions by BLaTM.** The eGFP-relative fluorescence ($\lambda_{exc}$ 485 nm, $\lambda_{em}$ 535nm) values for the BLaTM chimeras are shown. The mean and standard deviation of three independent experiments are shown (n = 3).

**a**

| | Sequence | ΔG$_{pred.}$ |
|---|---|---|
| **A** (wt) | FNRWFLTGMTVAGVVLLGSLFSR | -0.355 |
| **G** | FNRWFLTGMTV**G**GVVLLGSLFSR | 0.036 |
| **V** | FNRWFLTGMTV**V**GVVLLGSLFSR | -0.780 |
| **L** | FNRWFLTGMTV**L**GVVLLGSLFSR | -0.865 |
| **I** | FNRWFLTGMTV**I**GVVLLGSLFSR | -0.891 |
| **S** | FNRWFLTGMTV**S**GVVLLGSLFSR | 0.296 |
| **T** | FNRWFLTGMTV**T**GVVLLGSLFSR | 0.088 |
| **C** | FNRWFLTGMTV**C**GVVLLGSLFSR | -0.555 |
| **M** | FNRWFLTGMTV**M**GVVLLGSLFSR | -0.556 |
| **P** | FNRWFLTGMTV**P**GVVLLGSLFSR | 0.764 |
| **D** | FNRWFLTGMTV**D**GVVLLGSLFSR | 1.647 |
| **N** | FNRWFLTGMTV**N**GVVLLGSLFSR | 1.062 |
| **E** | FNRWFLTGMTV**E**GVVLLGSLFSR | 1.178 |
| **Q** | FNRWFLTGMTV**Q**GVVLLGSLFSR | 1.071 |
| **K** | FNRWFLTGMTV**K**GVVLLGSLFSR | 1.715 |
| **R** | FNRWFLTGMTV**R**GVVLLGSLFSR | 1.465 |
| **H** | FNRWFLTGMTV**H**GVVLLGSLFSR | 0.905 |
| **F** | FNRWFLTGMTV**F**GVVLLGSLFSR | -0.736 |
| **Y** | FNRWFLTGMTV**Y**GVVLLGSLFSR | 0.192 |
| **W** | FNRWFLTGMTV**W**GVVLLGSLFSR | -0.204 |

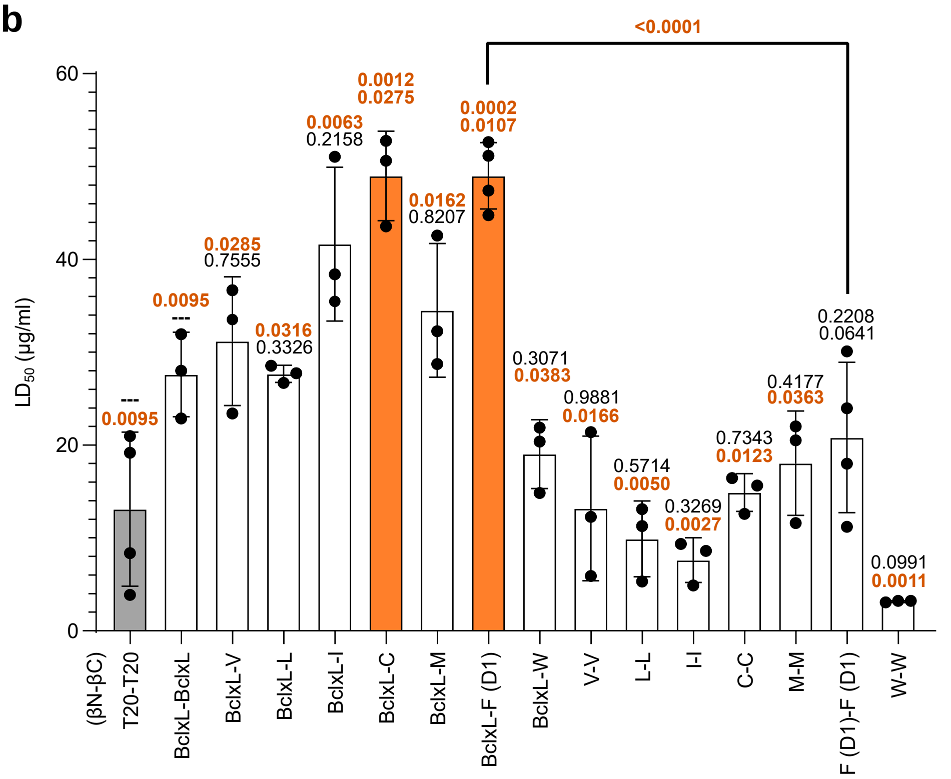


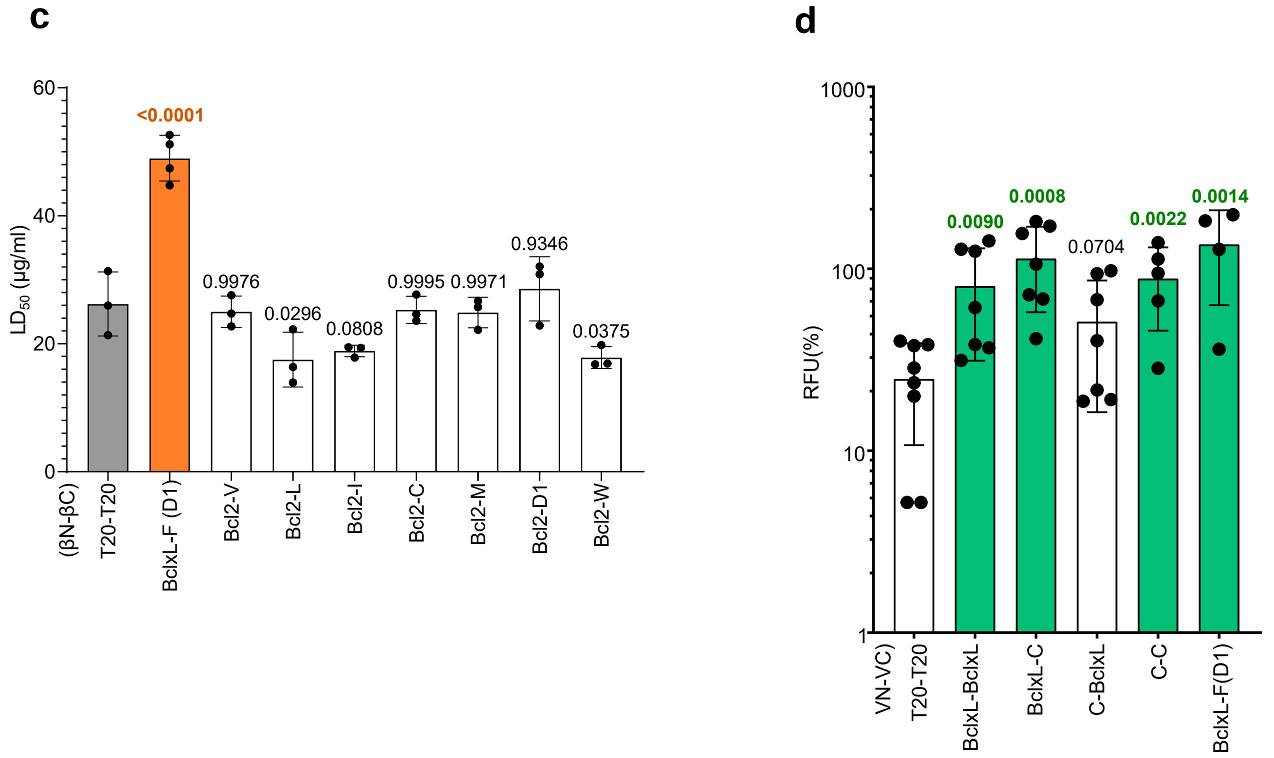


**Figure S6. Variations on a central position within the TMD of BclxL influence the insertion and interaction potential of the resulting segment.** The Ala in position 221 (position 12 in the TMD) was substituted, and the effect on the insertion into eukaryotic membranes was analyzed. The residue in position 12, the resulting TMD sequence, and the predicted insertion ΔGs (kcal/mol) calculated using the ΔG prediction server are shown (negative ΔGs in green, positive ΔGs in red). b. Substitutions that permitted insertion into the lipid bilayer were tested using the BLaTM assay to investigate their homo- and hetero-interaction (with the TMD of BclxL) potential. The indicated chimeras (βN-βC) were co-expressed in E. coli, and the resulting ampicillin LD50 was measured. The βN T20-βC T20 homodimer was used as a negative control (gray), and the βN GpA-βC GpA homodimer was used as a positive control to normalize values across experiments. The means and standard deviations of at least three independent experiments (n = 3) are shown. Values above the bars indicate the level of significance (ordinary one-way ANOVA test with Dunnett correction, p-value < 0.05) for the comparison with the T20 or BclxL TMD homo-interactions (top and bottom respectively). Additionally, the level of significance (ordinary one-way ANOVA test with Dunnett correction, p-value < 0.05) when comparing the interactions between the TMD of BclxL and D1 vs D1 and D1 is included. P-values < 0.05 are highlighted in bold orange letters, p-values > 0.05 are shown in black. c. Interaction of the BclxL TMD variants with the TMD of Bcl2. The indicated chimeras (βN-βC) were co-expressed in E. coli and the resulting ampicillin LD50 was measured. The means and standard deviations of at least three independent experiments are shown ($n \geq 3$). The individual value for each experiment is represented by a solid dot. The βN T20-βC T20 homodimer was used as a negative control (gray), and the βN GpA-βC GpA homodimer was used as a positive control and to normalize values across experiments (not included). The level of significance (p-value, ordinary one-way ANOVA test with Dunnett correction) when comparing the LD50 values of the indicated interactions vs T20 TMD homo-interactions is shown. d. BiFC was used to analyze the interaction potential of the BclxL TMD bearing the Ala12Cys substitution in eukaryotic membranes. The TMD included in each chimera (VN or VC) is indicated. The RFU mean and standard deviation of at least five independent experiments are shown. Solid dots represent the results of individual experiments. The TMD of T20 was used as negative control (gray). The TMD of GpA was used as a positive control and normalization value across experiments. An interaction (highlighted in green) was considered if the obtained RFU was significantly higher (two-tailed homoscedastic t-test, $p\text{-value} < 0.05$) than the negative control (gray bar). P-values are indicated above the corresponding bar (significantly higher in bold green). BclxL-F(D1) interaction was included as a control.

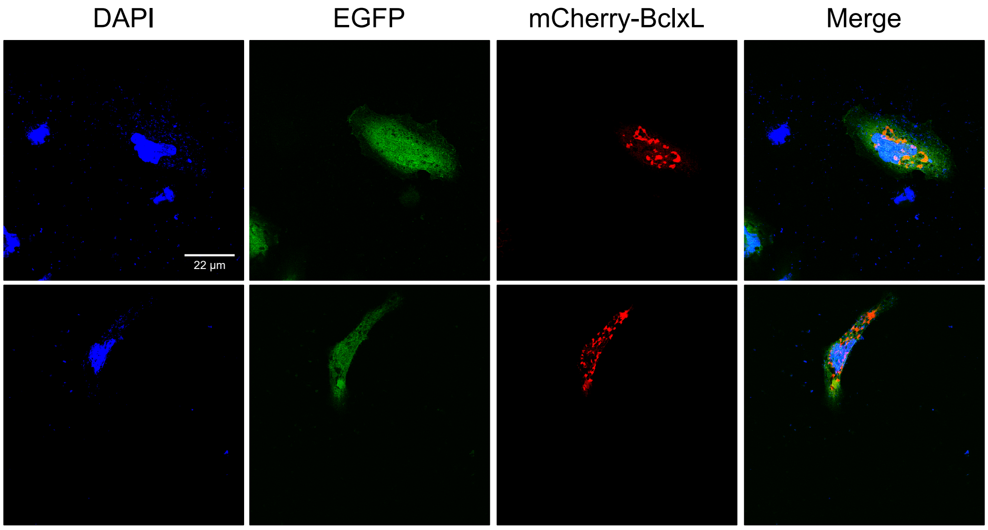


**Figure S7. Subcellular localization of the eGFP.** The eGFP was transfected into HeLa cells (green; $n = 3$) together with a mitochondrial marker (mCherry-BclxL, shown in red). DAPI staining is shown in blue. The right column of each panel shows the co-localization of both signals (yellow, visible only when the images are merged). No co-localization between eGFP and mCherry-BclxL was seen.

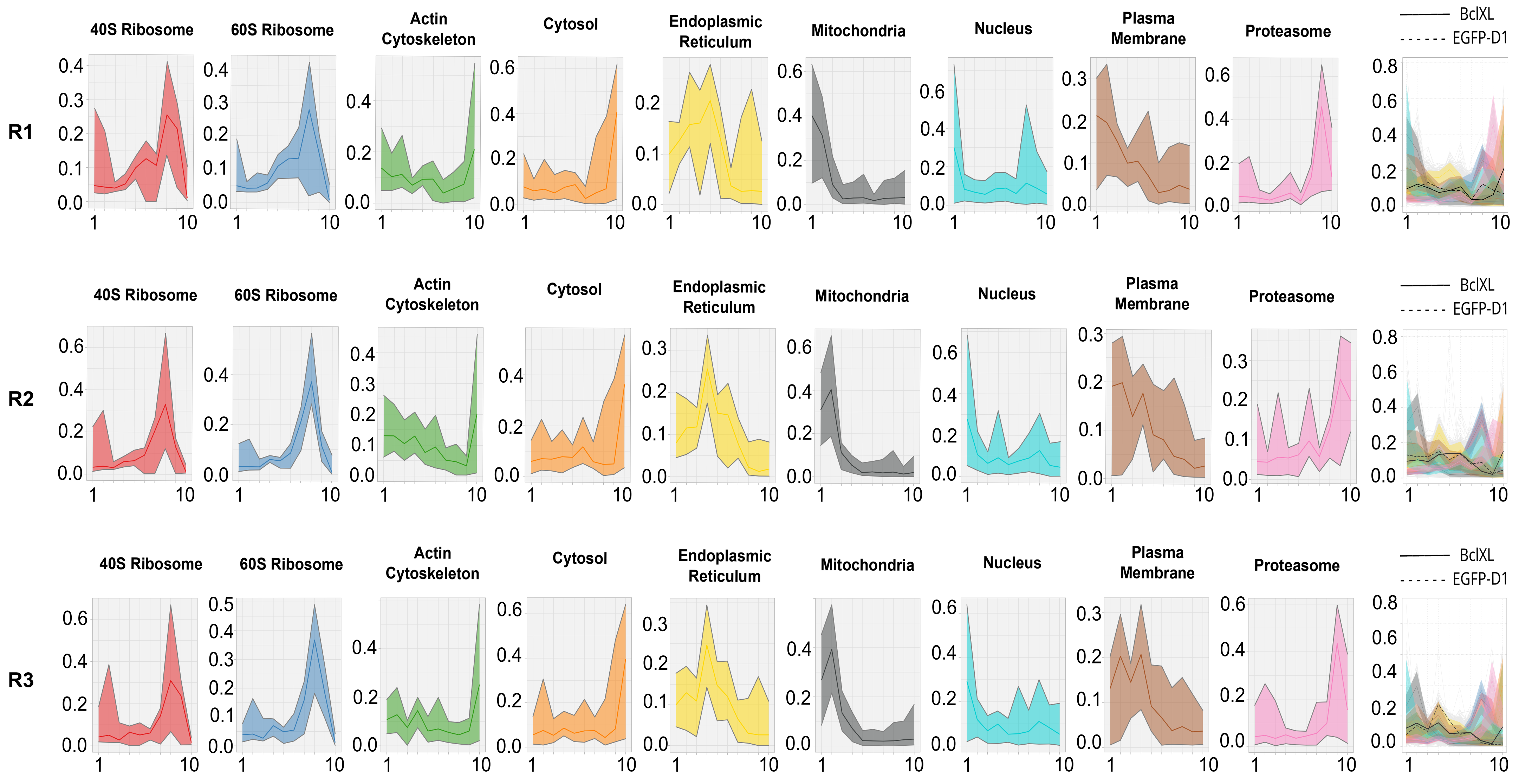


**Figure S8. Abundance profiles of organelle markers.** A series of differential ultracentrifugation steps was used to separate organelles and subcellular compartments. The proteins in each fraction were quantified by SWATH-MS analysis. The figure show the abundance profiles of the markers selected with the prolocGUI78 R package to identify organelles and subcellular compartments. The right panel shows the distribution of both BclxL (blue line) and eGFP-D1 (green dotted line) across fractions 1—10. The results of three independent experiments are shown (n = 3).

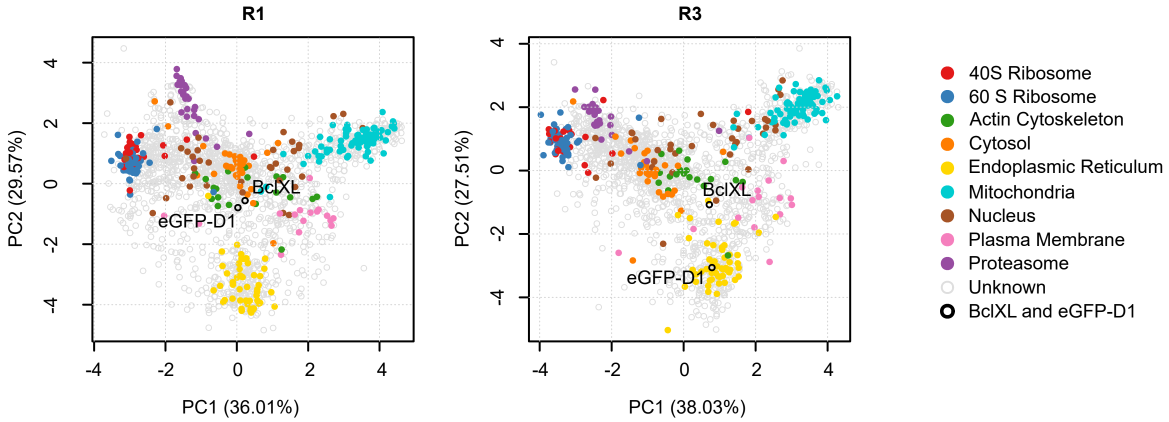


**Figure S9. Localization of BclxL and eGFP-D1 by differential centrifugation.** Principal Component Analysis (PCA) of the abundance profiles across fractions obtained by differential centrifugation. Each plot shows the 1962 proteins identified by SWATH-MS. The percentage in each axis represent the amount of total variability that PC1 and PC2 can explain. Organelle markers are colored. eGFP-D1 and BclxL indicated by white circles outlined in black, and the rest of the proteins (“other”) are indicated by in gray circles. The plot corresponding to replicate 2 was also included in figure 6.
